# Multicellular signaling in reverse cardiac remodeling revealed by integrated bulk and single-nucleus transcriptomics

**DOI:** 10.64898/2026.01.12.699040

**Authors:** Marco Steier, Ines Rivero-Garcia, Ricardo O. Ramirez Flores, Anushka Deshpande, Irsiya Aijaz, Julian Poetzsch, Sumra Nazir, Marina Talamini, Volker Ast, Katharina Mößinger, Matthias Dewenter, Benjamin Meder, Florian Leuschner, Manju Kumari, Johannes Backs, Ashraf Yusuf Rangrez, Julio Saez-Rodriguez, Norbert Frey

## Abstract

Heart failure results from maladaptive cardiac remodeling triggered by sustained biomechanical stress. Although mechanical unloading promotes reverse remodeling, recovery remains incomplete and its molecular basis poorly understood. Integrating functional assessment, bulk and single-nucleus transcriptomics in a reversible murine pressure-overload model, we characterized cell-type-specific recovery and persistent pathological programs. Unloading improved cardiac function and partially restored extracellular matrix and metabolic programs, while inflammatory and mitochondrial abnormalities persisted. Cardiomyocytes and endothelial cells largely regained homeostatic transcriptional states, whereas fibroblasts retained activated programs associated with residual remodeling. Multicellular factor analysis distinguished reversible and persistent transcriptional responses across cardiac cell types, while intercellular communication analysis nominated candidate recovery-associated signals. As one example, lymphatic endothelial Reelin and cardiomyocyte VLDLR recovered following unloading. Reelin attenuated cardiomyocyte hypertrophic responses in vitro, and circulating Reelin increased following transcatheter aortic valve replacement. Together, these findings reveal the heterogeneous cellular architecture of reverse remodeling and provide a framework for investigating novel recovery-associated pathways.

## 1. Introduction

Heart failure (HF) remains a leading cause of morbidity and mortality worldwide despite major therapeutic advances^1–3^. While contemporary pharmacological and device-based therapies improve survival and symptoms, they largely modulate disease progression rather than restore myocardial homeostasis, underscoring an incomplete understanding of the biological programs that govern both disease evolution and recovery^4–6^. Worsening of HF is driven by progressive adverse cardiac remodeling that impairs myocardial structure, function, and energetics. At the tissue level, HF is characterized by a complex constellation of pathological features, including cardiomyocyte hypertrophy and contractile dysfunction, reactivation of fetal gene programs, interstitial, bulk and perivascular fibrosis, chronic inflammation, microvascular rarefaction, and profound mitochondrial and metabolic reprogramming^7–10^. These alterations are not isolated events but reflect coordinated, system-level remodeling of the cardiac microenvironment. Increasing evidence indicates that pathological remodeling arises from dynamic interactions among cardiomyocytes, fibroblasts, endothelial cells, immune cells, and lymphatic endothelial cells, rather than from cardiomyocyte-autonomous processes alone^11^. Such multicellular coordination suggests that both disease progression and recovery are emergent properties of intercellular networks.

Importantly, cardiac remodeling is not strictly irreversible. In human, mechanical unloading most prominently achieved through left ventricular assist device (LVAD) support, can induce varying degrees of reverse remodeling in patients with advanced HF. Clinical studies have documented regression of cardiomyocyte hypertrophy, improvements in ventricular geometry, and reductions in myocardial fibrosis following unloading^12–14^. However, these changes are typically incomplete, and structural and molecular features of HF often persist at levels exceeding those observed in non-failing hearts^15^. Moreover, durable recovery after device explantation remains uncommon, suggesting that reverse remodeling may reflect incomplete restoration of cardiac homeostasis rather than a return to the non-failing state. Experimental models of pressure overload reversal mirror these clinical observations: relief of afterload improves systolic function and promotes regression of cardiomyocyte hypertrophy, yet residual fibrosis and architectural abnormalities frequently remain^16,17^. Together, these findings indicate that structural regression parallels functional recovery but remains incomplete, raising the possibility that persistent molecular and cellular programs actively constrain full reverse remodeling.

Transcriptomic profiling of failing human hearts and experimental models has revealed conserved activation of stress-responsive, proinflammatory, and metabolic pathways across cardiac cell types^18^. These studies have highlighted mitochondrial dysfunction, altered substrate utilization, and sustained inflammatory signaling as core features of HF pathophysiology. More recently, single-cell and single-nucleus RNA sequencing approaches have uncovered extensive cellular heterogeneity and dynamic cell-state transitions during cardiac remodeling^19–22^. These analyses have revealed persistent activation of fibroblast subsets, endothelial stress states, and immune-related transcriptional programs, even in contexts where global cardiac function improves, suggesting that incomplete recovery may reflect selective irreversibility at the cellular signaling or metabolic state level.

Despite these advances, important knowledge gaps remain. Many existing transcriptomic datasets capture static disease snapshots or late-stage human HF, limiting their ability to distinguish reversible from persistent remodeling programs. Moreover, while cell-intrinsic transcriptional changes have been well characterized, less is known about how multicellular programs shape recovery trajectories following unloading. Indeed, beyond cell-autonomous mechanisms, accumulating evidence implicates paracrine signaling in both pathological remodeling and repair. Fibroblast-derived extracellular matrix components, immune-derived cytokines, and endothelial-derived growth factors have all been shown to modulate cardiomyocyte growth, survival, and function^21,23^. In this context, cardiac lymphatic endothelial cells have emerged as active regulators of myocardial inflammation, metabolism, and tissue homeostasis through lymphoangiocrine signaling^24–26^. While lymphatic signaling has been implicated in cardiac repair after acute ischemic injury, its role in chronic HF and, in particular, in reverse remodeling following unloading remains largely unexplored.

Here, we addressed these gaps using a reversible murine model of pressure overload induced by O-ring aortic banding (ORAB) followed by surgical de-banding (DB). This approach enables controlled induction and subsequent relief of pressure overload within the same experimental system, providing an opportunity to distinguish reversible from persistent remodeling programs. By integrating detailed phenotypic characterization with bulk and single-nucleus RNA sequencing, multicellular factor analysis, and cell–cell communication inference, we systematically characterized the cell-type-specific responses, shared molecular programs, and intercellular signaling interactions associated with adverse and reverse cardiac remodeling. We further investigated lymphatic endothelial cell-derived Reelin as one candidate recovery-associated signal emerging from these analyses, combining myocardial signaling measurements, functional studies in rat and human cardiomyocytes, and circulating Reelin measurements in patients undergoing transcatheter aortic valve replacement (TAVR). Together, this integrated approach provides a framework for understanding the heterogeneous nature of myocardial recovery and identifying candidate pathways for future mechanistic investigation.

## 2. Results

### 2.1 Reversible pressure overload induces partial functional and structural cardiac recovery

To establish a controlled experimental framework for interrogating adverse and reverse cardiac remodeling, we employed a reversible murine model of pressure overload using O-ring aortic banding (ORAB) followed by surgical de-banding (DB). Eleven- to twelve-week-old mice were randomly assigned to four experimental groups: sham-operated controls euthanized two weeks after surgery, ORAB mice euthanized two weeks after banding, DB mice subjected to two weeks of ORAB followed by de-banding and an additional two-week recovery period, and time-matched sham controls euthanized four weeks after sham surgery to account for the extended duration of the DB protocol (**Fig. 1a**). Inclusion of a separate sham group for the DB cohort allowed us to control for potential time-dependent effects independent of pressure overload and unloading. As expected, ORAB induced a robust hypertrophic response and heart failure phenotype. Heart weights normalized to tibia length (HW/TL) were significantly increased in ORAB mice compared with sham controls, indicating pathological cardiac hypertrophy (**Fig. 1b**). De-banding led to a marked reduction in HW/TL, demonstrating significant regression of hypertrophy, although values remained modestly elevated compared with time-matched sham controls (**Fig. 1b**). These findings indicate that unloading triggers significant structural regression but does not fully restore baseline cardiac mass.

**Figure 1.**
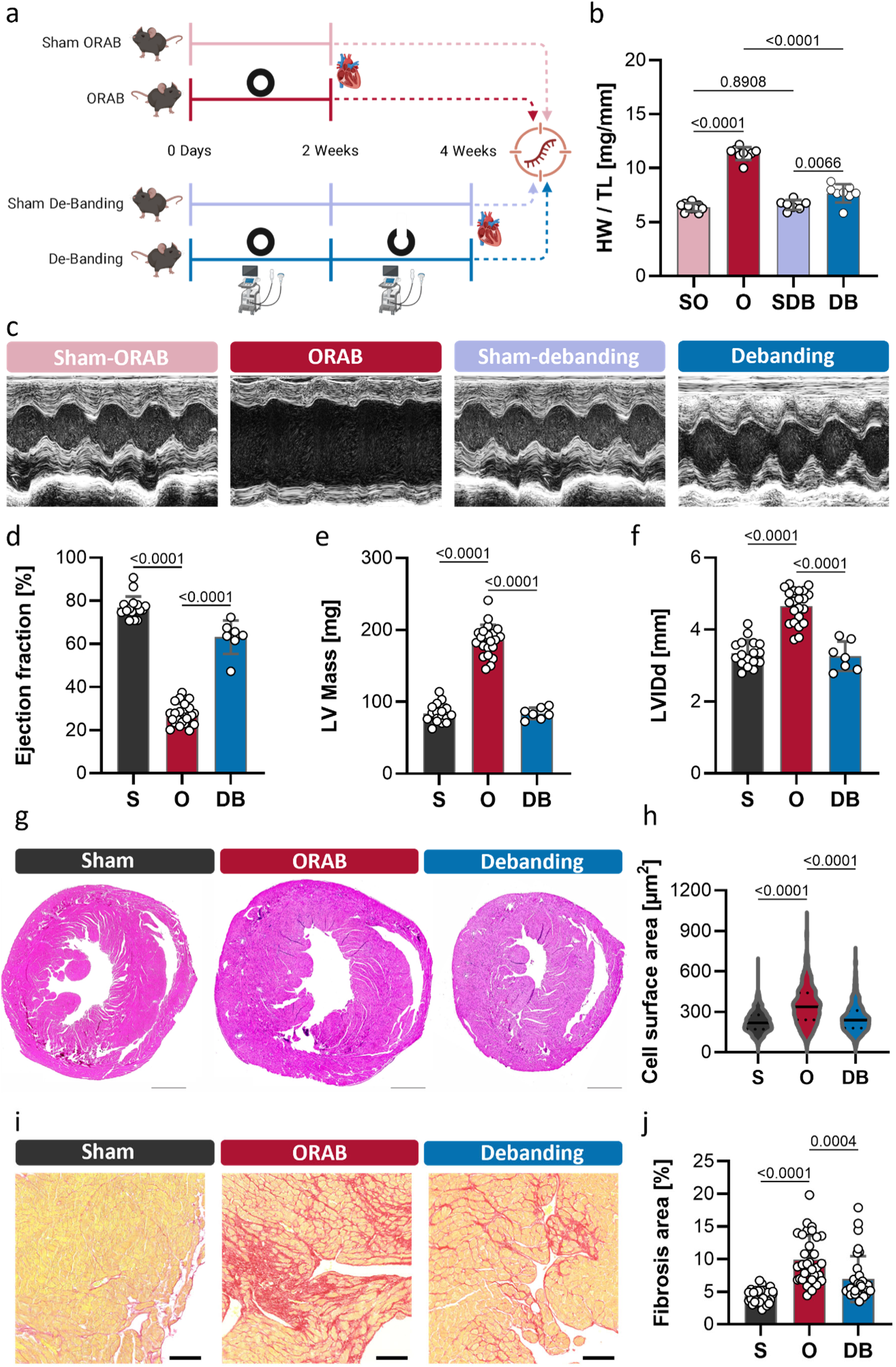
Reversal of pressure overload induces partial functional and structural cardiac recovery. **a)** Schematic of the experimental strategy. Eleven- to twelve-week-old mice were randomly assigned to sham control (SO), O-ring aortic banding (ORAB, or denoted as ‘O’ in the graph legend), de-banding (DB), or time-matched sham control for DB (SDB). ORAB mice were euthanized two weeks after banding. DB mice underwent two weeks of ORAB followed by surgical de-banding and an additional two-week recovery period. A separate sham group was included for DB to account for the extended experimental duration (this image is created at Biorender.com). **b)** Heart weight normalized to tibia length (HW/TL) showing increased cardiac hypertrophy after ORAB and significant, but incomplete, reduction after DB. n = 10 for SO, 9 for O, 7 for SDB and 8 for DB. **c)** Representative echocardiographic images illustrating severe cardiac dysfunction after ORAB and partial functional recovery following DB. **d)** Quantification of left ventricular ejection fraction (LVEF) demonstrating marked systolic impairment in ORAB mice and significant improvement after DB. n = 17 for sham (S), 22 for O, and 8 for DB. **e)** Left ventricular mass showing increased hypertrophy after ORAB and reduction following DB. n = 17 for sham (S), 22 for O, and 8 for DB. **f)** Left ventricular internal diameter at diastole, indicating pathological chamber dilation in ORAB mice and partial normalization after DB. n = 17 for sham (S), 22 for O, and 8 for DB. **g)** Representative hematoxylin and eosin-stained cardiac sections from sham, ORAB, and DB hearts, showing myocardial hypertrophy and structural remodeling after ORAB and partial regression after DB. **h)** Violin plot quantification of cardiomyocyte cross-sectional surface area demonstrating increased cell size after ORAB and significant reduction following DB. Cell surface area was measured from >200 cells per mouse heart, n = 3 mice per group. **i)** Representative fibrosis staining of sham, ORAB, and DB hearts, revealing extensive collagen deposition after ORAB and attenuated fibrosis after DB. **j)** Quantification of myocardial fibrosis showing severe fibrosis in ORAB hearts and partial but significant reduction after de-banding. Fibrotic area was quantified using 10 independent images acquired from comparable anatomical regions in three mice per group. Data are presented as mean ± SEM. Statistical significance was assessed using ordinary one-way ANOVA followed by Tukey’s multiple comparisons test. Exact p-values for the respective comparisons are indicated in the graph.

Echocardiographic assessment also revealed profound functional impairment in ORAB mice, characterized by reduced systolic performance and adverse ventricular remodeling. Representative M-mode images illustrated severe contractile dysfunction after pressure overload, which was partially rescued following de-banding (**Fig. 1c**). Quantitative analysis confirmed a marked reduction in left ventricular ejection fraction (LVEF) and fractional shortening (LVFS) in ORAB mice, consistent with overt heart failure, whereas DB mice exhibited a significant improvement in LVEF and LVFS, though not complete normalization to sham levels (**Fig. 1d** and **Fig. S1a, b**). In parallel, left ventricular mass was substantially increased after ORAB and significantly reduced after de-banding (**Fig. 1e** and **Fig. S1c**). Similarly, left ventricular internal diameter at diastole and systole were increased in ORAB mice, reflecting pathological chamber dilation, and was significantly reduced following unloading (**Fig. 1f** and **Fig. S1d, e**). Together, these data demonstrate that de-banding promotes partial recovery of cardiac function and geometry.

To complement these phenotypic and functional assessments, we evaluated expression of canonical fetal and stress-associated genes by quantitative PCR. Transcripts encoding atrial natriuretic peptide (*Nppa*) and brain natriuretic peptide (*Nppb*) were markedly upregulated in ORAB hearts, consistent with pathological hypertrophy and cardiac stress. De-banding significantly reduced expression of all three genes, although levels did not fully normalize to sham controls (**Fig. S1f**). Notably, all phenotypic and functional parameters, including HW/TL, echocardiographic measurements, representative M-mode images, and gene expression were indistinguishable between the two sham groups euthanized at two or four weeks after surgery (**Fig. 1b, c** and **Fig. S1a-f**). Based on this equivalence, data from both sham cohorts were combined for echocardiographic analyses shown in **Fig. 1d-f** and were subsequently pooled for all downstream structural, molecular, and transcriptomic analyses.

Histological analyses further corroborated these findings at the cellular and tissue levels. Hematoxylin and eosin-stained cardiac sections revealed marked myocardial thickening and architectural distortion in ORAB hearts compared with sham controls, whereas DB hearts exhibited visibly reduced wall thickness and improved myocardial organization, albeit without complete normalization (**Fig. 1g**). Quantification of cardiomyocyte cross-sectional surface area showed a pronounced increase in ORAB mice, indicative of pathological hypertrophy. De-banding resulted in a significant reduction in cardiomyocyte size, yet cardiomyocytes remained larger than those in sham hearts (**Fig. 1h**), consistent with incomplete regression of hypertrophy. Assessment of myocardial fibrosis revealed extensive interstitial and perivascular collagen deposition in ORAB hearts, again reflecting advanced pathological remodeling. In contrast, DB hearts displayed a significant reduction in fibrotic burden, although residual fibrosis persisted relative to sham controls (**Fig. 1i**). Quantitative analysis confirmed a robust increase in fibrosis following ORAB and a partial but significant reversal after unloading (**Fig. 1j**).

Collectively, these phenotypic, functional, molecular and histological data establish that reversal of pressure overload through de-banding induces significant improvement in cardiac structure and function but fails to fully restore a non-diseased state. This model therefore provides a robust platform to dissect the molecular and multicellular programs that govern incomplete reverse remodeling. While these phenotypic, functional, and molecular assessments establish that unloading induces substantial yet incomplete cardiac recovery, they do not reveal the molecular programs that enable recovery or constrain its extent. Thus, to define the transcriptional mechanisms underlying this intermediate remodeling state, we next performed bulk RNA sequencing (RNA-seq) followed by single-nucleus RNA sequencing (snRNA-seq).

### 2.2 The cardiac transcriptome is partially recovered after pressure overload relief

Unsupervised analysis of the bulk RNA-seq data by Principal Component Analysis (PCA) showed clear separation between the three groups (variance explained by PC1 = 63%, **Fig. S2a, b**), with a damage-to-recovery gradient captured by PC1 (ANOVA Bonferroni-corrected p-value = 1.57e-06) (**Fig. 2a, b**). To validate the existence of this gradient, we calculated the Euclidean distance between all pairs of samples using the normalised expression of all genes with more than 10 counts in at least 3 samples. The results agreed with the gradient observed in PC1. The ORAB-Control comparisons show the largest distance, followed by the distances between ORAB and DB, and DB and Controls (Kruskal-Wallis rank sum test p-value < 2.2e-16) (**Fig. 2c**). Because these two unsupervised analyses showed that the two control groups (Sham-ORAB and Sham-DB) are virtually identical to each other, they remained grouped as Controls for further analyses (**Fig. S2c**).

**Figure 2.**
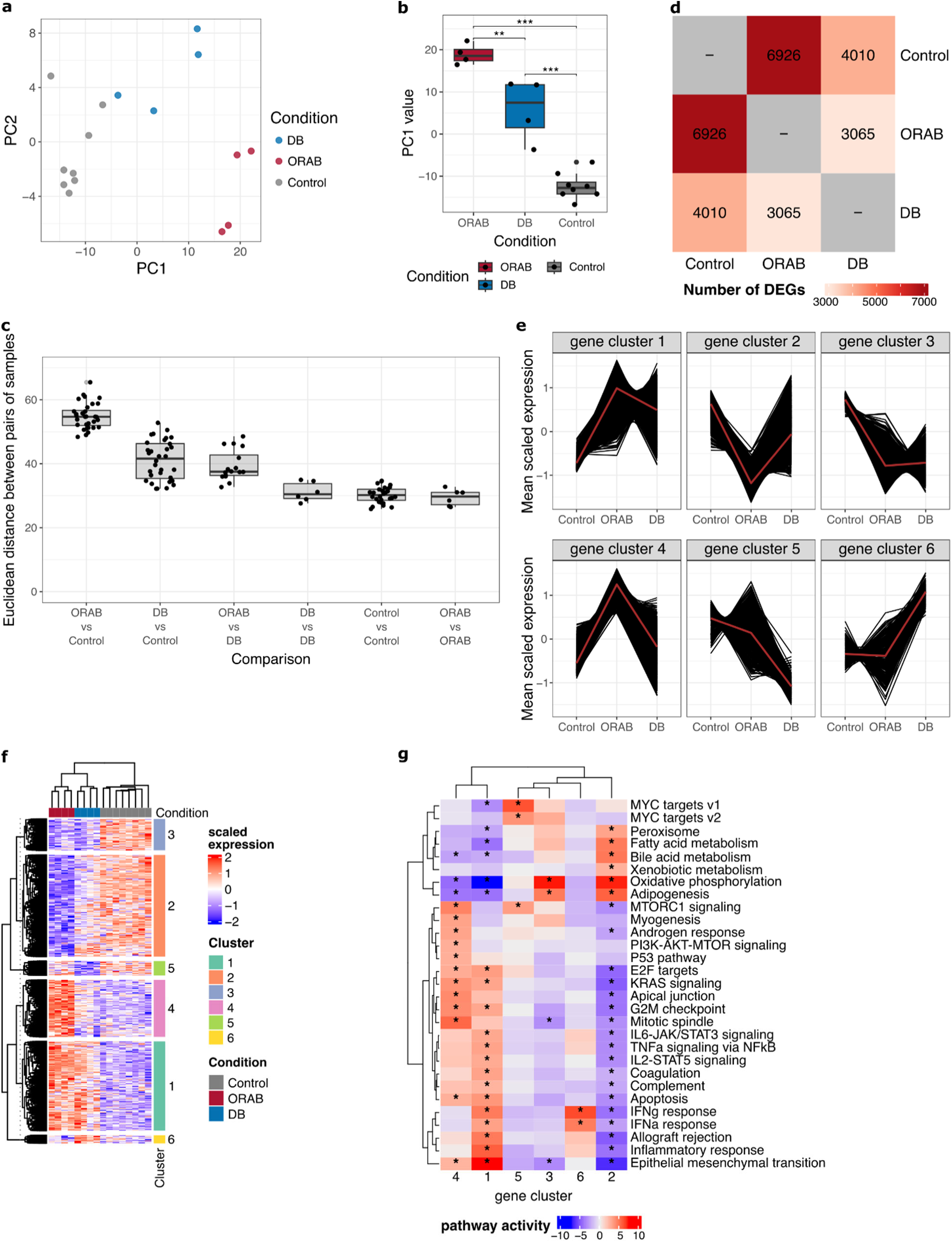
Bulk RNA-seq identified a partial transcriptional recovery 2 weeks after DB. **a)** PCA visualisation of the bulk RNA-seq. Each point in the scatter plot represents a biological sample. **b)** Boxplot of the PC1 scores. Each dot indicates a biological replicate. **c)** Boxplot of the euclidean distances between pairs of samples calculated based on their bulk mRNA expression levels. Each point represents the euclidean distance between a pair of samples. **d)** Heatmap of the number of DEGs detected between conditions. **e)** Mean scaled expression values of DEGs grouped by k-means clustering across conditions. Each line represents the average of four biological replicates per time-point. Black lines indicate the average expression of individual genes across biological replicates (n = 8 for Control, n = 4 for DB and n = 4 for ORAB), and the red line indicates the average expression of all genes in the cluster. **f)** Heatmap of the scaled expression of the differentially expressed genes grouped by cluster. **g)** Heatmap of the activity scores for the top pathways overrepresented in each gene cluster. Asterisks indicate FDR-adjusted p-values < 0.05.

To identify from the gene expression processes that showed variability between ORAB, DB and Controls, we calculated the differentially expressed genes (DEGs) for all possible contrasts. The number of DEGs further supports the damage-to-recovery gradient, as 6,926 DEGs are detected when comparing ORAB vs. Controls, but only 3,065 and 4,010 DEGs are detected when comparing ORAB to DB and DB to Controls, respectively (**Fig. 2d**, **Fig. S2d** and **Sup. File 1**). Pearson correlation of the gene expression changes (FDR < 0.05 in at least one comparison) show that the gene expression differences in ORAB vs Control are very similar to those in ORAB vs DB (Pearson r = 0.85) and to those in DB vs Control (Pearson r = 0.90). Interestingly, this correlation is lower when the log2FC of DEGs in ORAB vs DB are compared to the log2FC of DEGs in DB vs Control (Pearson r = 0.47). Adding support to the damage-recovery gradient, these results suggest that some biological processes characteristic of Controls are not fully reversed in DB (**Fig. S2e**). To explore the reversibility of the adverse remodelling transcriptome in more detail, we performed hierarchical clustering of the DEGs (genes with FDR < 0.05 in at least one condition) and identified six clusters with different behaviours across the experimental timeline (**Fig. 2e, f** and **Sup. File 2**). Two of the clusters (gene clusters 1 and 4) capture partially reversible upregulation, as the expression of these genes was sharply increased after ORAB, and partially reversed to Control levels after DB. Gene cluster 1 is characterized by classical tissue remodelling and fibroblast activation functions, including epithelial-mesenchymal transition, TNFa signaling via NFkB and G2/M checkpoints (**Fig. 2g** and **Sup. File 3**). Genes in this cluster include collagen subunits (i.e. *Col1a1*, *Col1a2*, *Col3a1*), matrix metalloproteases (i.e. *Mmp2*, *Mmp14*, *Adamts2*) fibroblast activation markers (i.e. *Postn*, *Tgfb2, Tgfbr1*) and G2/M cell cycle regulators (i.e. *Ccnb1*, *Cdk1* and *Mcm5*) (**Fig. S2f**). While some of these pathways are also enriched in Gene cluster 4, gene cluster 4 presents significant upregulation in metabolic sensing and rewiring and DNA damage checkpoints (**Fig. 2g** and **Sup. File 3**). Its genes include metabolic sensors (i.e. *Rragc*, *Prkab2*, *Akt3*), glucose metabolism (i.e. *Pgk1*, *Hk1*, *Ldha*), G1/S replication checkpoint genes (i.e. *Ccne1*, *Cdkn1a*, *E2f7*) and mediators of p53-induced apoptosis (i.e. *Perp*, *Bax*, *Bbc3*) (**Fig. S2f**).

One of the clusters (gene cluster 2) captures partially reversible downregulation, as the expression of these genes was sharply reduced after ORAB and partially recovered after DB. Gene cluster 2 is associated with fatty acid metabolism particularly in the peroxisome (**Fig. 2g** and **Sup. File 3**). The peroxisomal biogenesis genes *Pex6*, *Pex7* and *Pex12*; the fatty acid metabolism regulators *Pparg*, *Ppargc1a* and *Ppara*; and the peroxisomal fatty acid metabolism enzymes *Acox1, Acot1* and *Acot6* are part of this cluster (**Fig. S2f**).

Interestingly, the other three clusters captured irreversible gene expression changes. Two of them (gene clusters 3 and 5) capture irreversible downregulation i.e. processes that are downregulated after ORAB and not recovered during DB. However, the time dynamics of these two clusters follow different trends. Gene cluster 3 comprises gene expression changes that are switched off during ORAB, and remain at a similar level after DB. Genes in this cluster are enriched in mitochondrial oxidative phosphorylation (**Fig. 2g** and **Sup. File 3**), mitochondrial electron transport chain components and assembly factors (i.e. *Ndufa1*, *Cox5b*, *Atp5po*), members of the mitochondrial transcriptional machinery (i.e. *Tfam*, *Tefm*, *Mtpap*) and regulators of mitochondrial dynamics (i.e. *Opa1*, *Mtfr1*, *Mfn2*) (**Fig. S2f**). Gene cluster 5 on the other hand captures a gradual downregulation of gene expression. Transcriptional targets of MYC (i.e. *Rpl19*, *Nolc1*, *Ldhb*) and genes related with MTORC1-signaling (i.e. *Akt1*, *Sirt2*, *Insr*) are overrepresented in this cluster (**Fig. 2g**, **Fig. S2f** and **Sup. File 3**).

Lastly, gene cluster 6 comprises genes whose expression is not significantly changed between ORAB and Controls, but increased after DB. This cluster is associated with IFNα and IFNγ signaling (**Fig. 2g** and **Sup. File 3**). The interleukin receptors *Il2rb* and *Il11ra1*; the JAK/STAT transcription factor *Stat1*; and the interferon-induced genes *Ifit1 and Mx2* belong to this cluster (**Fig. S2f**).

Overall, bulk RNA-seq analysis suggested that DB may be an intermediate phenotype between ORAB and Controls at the gene expression level. While some pathological changes such as collagen deposition, fibroblast activation and peroxisomal fatty acid metabolism may partially revert two weeks after DB, other pathological pathways such as inflammation and impaired oxidative mitochondrial metabolism remain de-regulated two weeks after DB.

### 2.3 Cellular landscape of adverse and reverse cardiac remodeling

To understand the extent to which this partial reverse remodelling is due to irreversible changes in the proportion of cell types or to changes in gene expression, we performed snRNA-seq of the ORAB, DB and Controls, following the same experimental design as before. A total of 174,391 nuclei passed quality control (**Fig. S3** and **Table S1**) and integration showed homogeneous mix of distinct batches while preserving biological differences between conditions (**Fig. S4**). Clustering of the integrated snRNA-seq data identified 27 clusters that were annotated to 9 major cell types based on the expression of canonical, cell type markers as described in the meta-analysis of single-cell HF study^21^ and CellTypist heart models^27^. These 9 major cell types comprise cardiomyocytes (CM) (*Ryr2*^+^, *Fhl2*^+^, *Tcap*^+^, *Actn2*^+^), endothelial cells (*Kdr*^+^, *Cdh5*^+^, *Eng*^+^, *Tie1*^+^), fibroblasts (*Col1a1*^+^, *Pdgfra*^+^, *Col1a2*^+^, *Col3a1*^+^), lymphatic endothelial cells (LECs) (*Lyve1*^+^, *Prox1*^+^, *Flt4*^+^, *Pdpn*^+^), lymphoid cells (*Ighm*^+^, *Ighd*^+^, *Pax5*^+^, *Stat4*^+^), myeloid cells (*Adgre1*^+^, *Csf1r*^+^, *C1qa*^+^, *C1qb*^+^), neuronal cells (*Plp1*^+^, *Lgi4*^+^, *Cadm2*^+^, *Xkr4*^+^), pericytes (*Rgs5*^+^, *Abcc9*^+^, *Kcnj8*^+^, *Pdgfrb*^+^) and vascular smooth muscle cells (vSMCs) (*Acta2*^+^, *Myh11*^+^, *Tagln*^+^, *Myl9*^+^) (**Fig. 3a, b**). Two small clusters were labelled as unknown, as they expressed markers of several cell types and we weren’t able to confidently annotate them. These two clusters were excluded from further analysis (**Fig. S5** and **Sup. File 4**). Additionally, the neuronal cell population was excluded from further analyses due to the low number of nuclei detected per sample (**Fig. S6a-c**).

**Figure 3.**
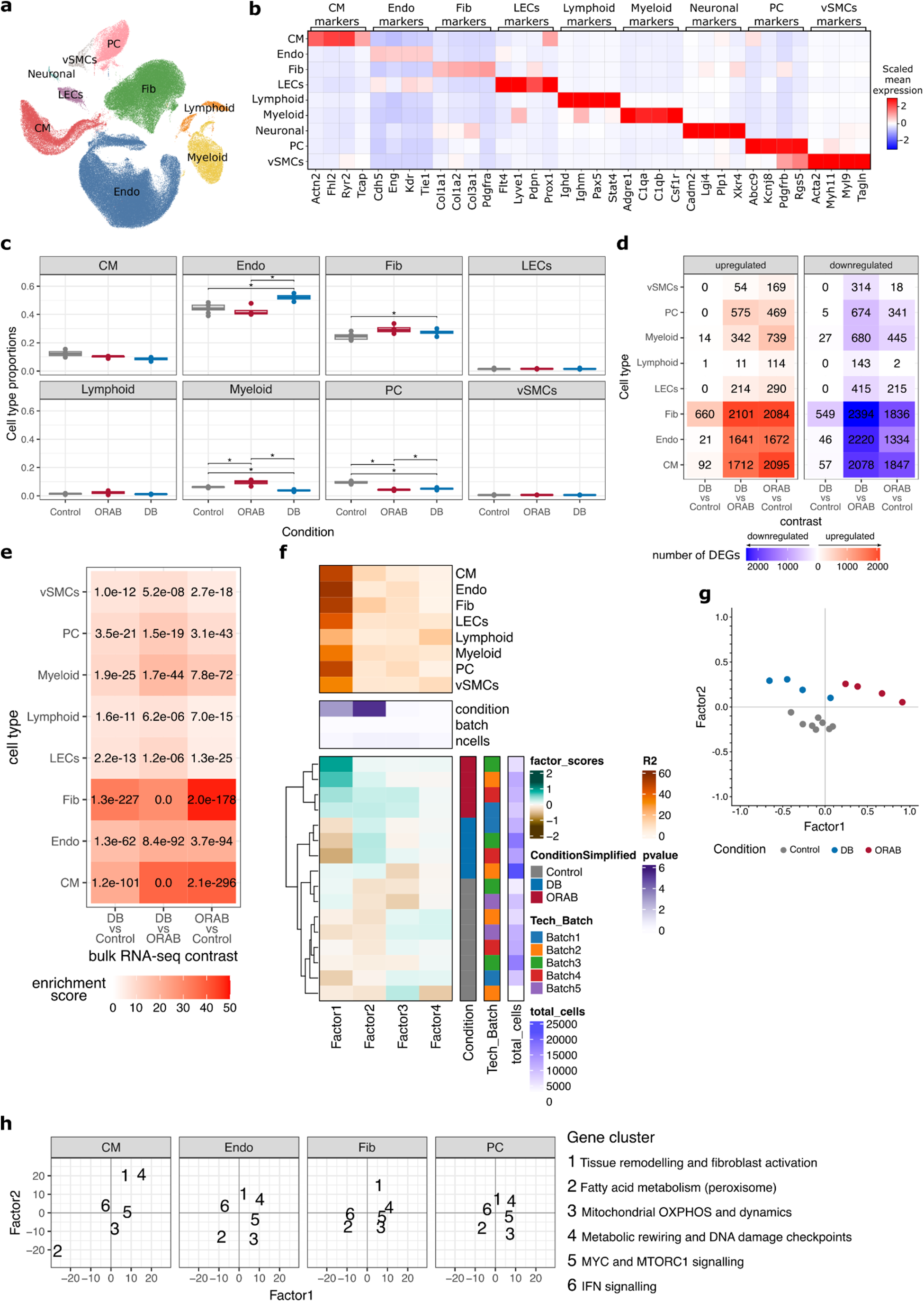
Changes in cell type composition and coordination during adverse and reverse remodelling. **a)** UMAP visualisation of the snRNA-seq, colored by major cell types. **b)** Heatmap of the scaled and log-normalised expression of canonical marker genes for the major cell types identified in the snRNA-seq. c) Boxplots of the cell type proportions present in each sample. Each dot represents one biological replicate. Asterisks indicate Tukey HSD adj. p-values < 0.05 for contrasts with ANOVAs adj. p-values < 0.05. **d)** Heatmap of the number of DEGs detected per cell type for the DB vs. Control, DB. vs ORAB and ORAB vs. Control contrasts. Red indicates upregulated genes (fdr < 0.05 and log2FC > 0) and blue indicates downregulated genes (fdr < 0.05 and log2FC < 0). **e)** Heatmap of the enrichment scores of cell type-specific log2FCs in the bulk RNA-seq DEGs. The FDR adjusted p-values of each enrichment are written in the corresponding heatmap cell. **f)** Heatmap of the Multicellular Factor Analysis model factor scores, factor association p-values and R2 coefficients. **g)** Scatterplot of the sample weights for multicellular factors 1 and 2. **h)** Scatterplot of the enrichment scores of the gene clusters identified in bulk in the cell type-gene loadings of multicellular factors 1 and 2. Only the main contributing cells are shown.

Due to availability of cell type resolution, we conducted a compositional analysis using ANOVA to identify if any of the major cell types change in their abundance upon ORAB or DB. We identified significant differences in the proportion of endothelial cells (ANOVA adj. p-value = 6.42e-04), pericytes (ANOVA adj. p-value = 1.80e-07), myeloid cells (ANOVA adj. p-value = 1.16e-04) and fibroblasts (ANOVA adj. p-value = 0.020) across conditions (**Fig. 3c** and **Table S2**). The proportion of endothelial cells increased as a consequence of DB (ORAB vs. DB Tukey HSD adj. p-value = 4.18e-04), reaching levels higher than what was observed in the Controls (DB vs. Control Tukey HSD adj. p-value = 5.96e-04). These results are consistent with the reported increase in the coronary microvasculature network seen in human samples post-LVAD implantation^28^. With respect to pericytes, their abundance decreased after ORAB (ORAB vs Control Tukey HSD adj. p-value = 2.26e-08), as previously reported^29^. Despite an increase after DB (ORAB vs DB Tukey HSD adj. P-value = 1.56e-03), they did not recover Control levels (DB vs Control Tukey HSD adj. p-value = 8.47e-06). As for myeloid cells, we observed an increased abundance in ORAB hearts (ORAB vs Control Tukey HSD adj. p-value = 1.91e-03 and ORAB vs DB Tukey HSD adj. p-value = 1.95e-05), which may recapitulate the known macrophage infiltration that takes place during early ischemic remodelling^23^. Regarding fibroblasts, we observed a significant increase in DB compared to Controls (Tukey HSD adj. p-value = 0.010). Lastly, we observed a decreasing tendency in the abundance of CMs, although this was not significant (ANOVA adj. p-value = 0.076). Overall, these results suggest that predominant changes in the myocardial vasculature might occur as a consequence of DB, with increased endothelial cell abundance but irreversible pericyte loss.

Next, we explored how adverse and reverse remodelling affected the gene expression profile of each major cell type. For this, we generated sample and cell type-specific pseudobulks by averaging the expression of individual cells of the same type and sample (**Fig. S6**), and calculated the differentially expressed genes between conditions. Overall, the cell types showing the largest numbers of DEGs were fibroblasts, CMs and Endothelial cells (**Fig. 3d** and **Sup. File 5**). Differently to what we observed in the bulk RNA-seq, the number of DEGs detected in ORAB vs. Control and ORAB vs. DB was very similar, while few DEGs were detected in the DB vs. Control contrast (**Fig. 3d** and **Fig. S7a**). However, enrichment of the cell type-specific log2FCs in the bulk log2FCs showed an agreement of changes in gene expression for all cell types and contrasts, supporting that the directionality of gene expression changes is conserved between technologies, despite differences in statistical significance (**Fig. 3e**). Inspection of the cell type-specific Euclidean distances between the pseudobulks showed that the damage-to-recovery gradient was only observable in fibroblasts (**Fig. S7b**).

Because individual cell types do not function alone but rather are exposed to the same microenvironment, we fitted a Multicellular Factor Analysis model^30^ to identify multicellular programs capturing coordination between different cell types. This unsupervised analysis identified 2 factors significantly associated with the experimental groups (**Fig. 3f**). Factor 1 (ANOVA adj. p-value = 1.07e-03) displayed coordinated variability across all cell types (mean R2 = 0.42, max. R2 = 0.55, min R2 = 0.22) (**Table S3**) and captured reversible multicellular coordination, as its loadings are increased in ORAB samples compared to DB and Controls. Factor 2 (ANOVA adj. p-value = 5.05e-06) captured a smaller variability, being most important in Fibroblasts (R2 = 0.15) and CMs (R2 = 0.12) (**Table S3**). Factor2 captured irreversible multicellular coordination, as its loadings are higher in ORAB and DB samples compared to Controls (**Fig. 3g**).

To understand how these factors relate to the gene clusters identified in the bulk RNA-seq, we enriched the cluster genes in the cell type gene loadings for each factor and cell type. The results show that both factors captured the gene expression changes described by the bulk clusters in a cell type-dependent fashion (**Fig. 3h** and **Fig. S8**). For example, the fibrosis (gene cluster 1) and metabolic rewiring and DNA damage checkpoints (gene cluster 4) were mostly activated in CMs and fibroblasts of ORAB hearts. On the other hand, peroxisomal fatty acid metabolism (gene cluster 2) and mitochondrial OXPHOS and dynamics (gene cluster 3) were mostly activated in CMs, endothelial cells and pericytes of Control hearts.

Overall, the snRNA-seq identified an increased myeloid infiltration in ORAB samples and an increase in the coronary vasculature because of DB. The cell type-specific DEGs revealed that most expression changes are derived from CMs, endothelial cells and fibroblasts, while the damage-to-recovery gradient could only be detected in fibroblasts. Lastly, the multicellular programs identified two Factors across cell types that captured the reversible and irreversible gene expression changes and provided insights into cell type-specific contributions.

### 2.4 De-banding restores the homeostatic cardiomyocyte state

To explore whether the transcriptomic changes between ORAB, DB and Controls arise from global changes in a cell type expression or from specific populations, we performed sub-clustering of the major cell types to identify cell states. We performed this analysis on CMs, endothelial cells and fibroblasts, as they showed the largest differences between conditions.

Sub-clustering of CMs identified 10 clusters. Three of them (clusters 0, 1 and 2) comprised almost 90% of the total CM nuclei, and two of them showed compositional differences between conditions (**Fig. 4a, b** and **Table S4**). CM cluster 0 was virtually depleted from ORAB nuclei and equally abundant in Control and DB hearts (ANOVA adj. p-value = 9.25e-06). This cluster resembled the vCM1 population identified by Litvinukova *et al*^31^, characterised by being the most abundant ventricular CMs in the healthy human left ventricle^31^ (**Fig. 4c**). These CMs expressed marker genes characteristic of homeostatic CM functions, such mitochondrial fatty acid oxidation (i.e. *Ppara*, *Ppargc1b*, *Cpt1a*), electrical coupling (i.e. *Gja1*), mechanical coupling (i.e. *Dsg2*, *Dsc2*) and regulation of ion levels (i.e. *Kcnj2* and *Cacnb2*) (**Fig. 4d**). CM cluster 2 was almost uniquely composed of ORAB CMs (ANOVA adj. p-value = 7.13e-03). CM cluster 2 showed increased expression of genes associated with cardiac stress, such as the fetal CM markers (i.e. *Nppa*, *Nppb* and *Myh7),* genes involved in the metabolic glycolytic shift (i.e. *Hif1a*, *Pfkp* and *Gck*) and genes involved in fibrosis (i.e. *Tgfb2*, *Ccn1*, *Ccn2*) (**Fig. 4d, e** and **Sup. File 6**). Additionally, the *Myh7*/*Myh6* expression ratio in these CMs is significantly higher than in the basal CMs captured in cluster 0 (ANOVA adj. p-value = 0.002) (**Fig. 4f**).

**Figure 4.**
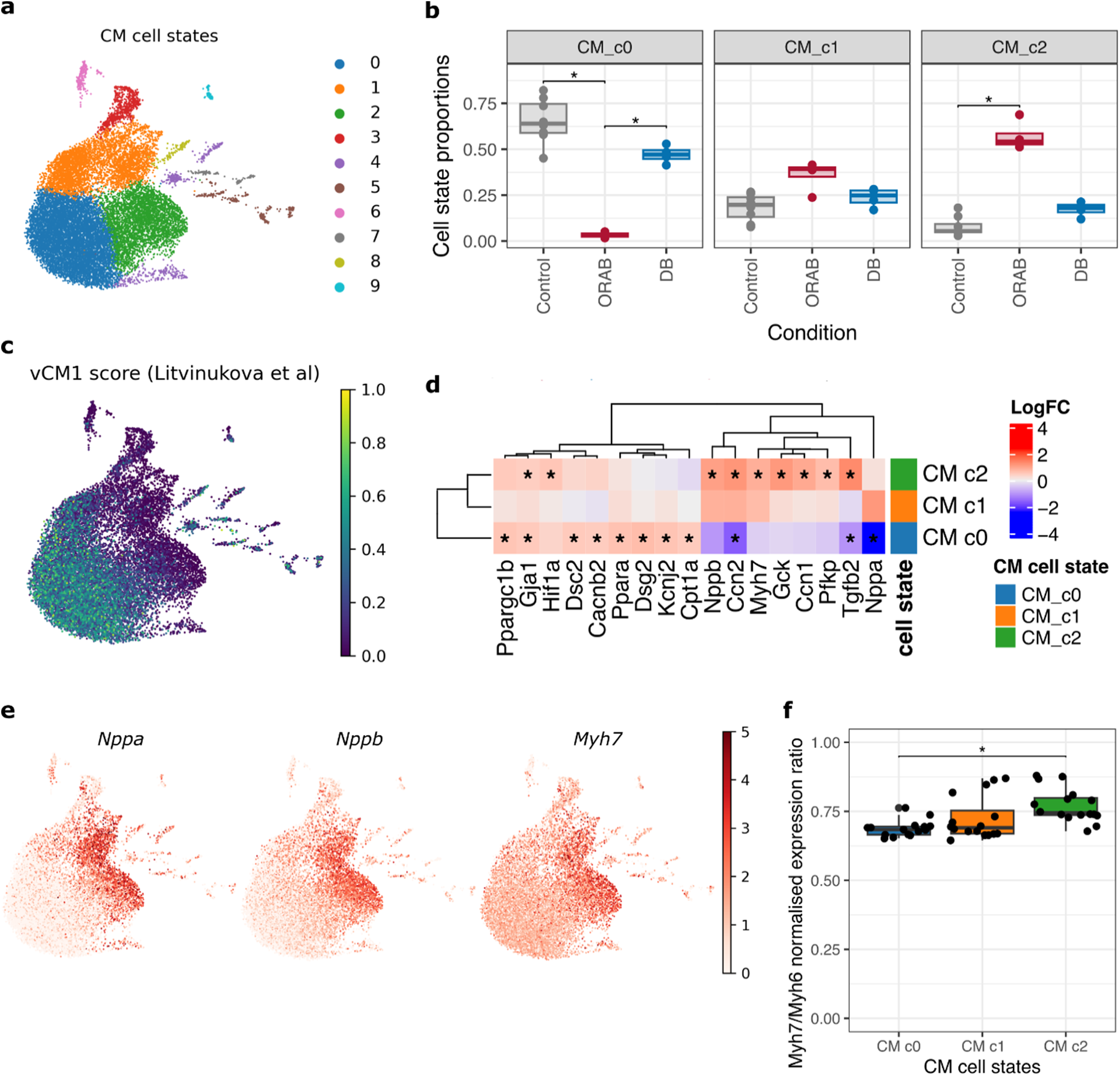
Changes in CM cell states during adverse and reverse remodelling. **a)** UMAP visualisation of the CM cell states. **b)** Boxplots of the cell type proportions for the CM cell states representing > 5% of all CMs. Each dot represents one biological replicate. Asterisks represent Tukey HSD adj. p-value < 0.05, for contrasts with ANOVA adj. P-value < 0.05. **c)** UMAP visualisation of the CMs, colored by the CellTypist score of similarity with the vCM1 population described by Litvinukova et al.32. **d)** Heatmap of the LogFC of selected marker genes in the main CM cell states. The logFC were calculated using all other CMs as reference. Asterisks indicate FDR < 0.05. **e)** UMAP visualisation of the CMs, colored by their log-normalised expression of the fetal marker genes Nppa, Nppb and Myh7. f) Boxplot of the ratio of Myh7/Myh6 normalised expression in the main CM cell states. The asterisks indicate a Tukey HSD adj. P-value < 0.05.

Regarding endothelial cells, we identified 8 clusters, among which some resembled the capillary (clusters 0, 2 and 6), arterial (cluster 3 and 7) and venous (clusters 1 and 4) ventricular endothelial cells described by Litvinukova *et al*.^31^ (**Fig. S9a, b**). Four of the clusters showed differences in composition across conditions, with two clusters being depleted in ORAB and two clusters being increased in ORAB (**Fig. S9c** and **Table S5**). Endothelial clusters 0 and 1 were depleted in ORAB samples (cluster 0 ANOVA p.adj = 7.44e-06 and cluster 1 ANOVA p.adj = 7.44e-06). Marker genes for endothelial cluster 0 include genes involved in cellular junctions (i.e., *Cdh5*, *Tip1*, *Jam2*) and fatty acid metabolism (i.e. *Lpl*, *Fh1*, *Cd36*) (**Fig. S9d**). Marker genes for endothelial cluster 1 include venous markers (i.e. *Ephb4*, *Nr2f2*) and immunosurveillance (i.e. *Il1r1*, *Icam1*, *Il4ra*) (**Fig. S9d**). On the other hand, endothelial clusters 2 and 4 are more abundant in ORAB than in DB or Control samples (cluster 2 ANOVA adj. p-value = 4.34e-04 and cluster 4 ANOVA adj. p-value = 3.54e-04). Marker genes for endothelial cluster 2 include classic tissue remodelers (i.e. *Nfkb1*, *Nfkb2*, Rel, Relb, Tgfb1), mediators of inflammation (i.e., Ifngr1, Ifnar1, Il16, Csf1) and proliferation (i.e. *Mki67*, *Cdk6*, *Ccnd1*) (**Fig. S9d** and **Sup. File 7**). Marker genes for endothelial cluster 4 include hypoxia-induced stress mediators (i.e. *Apln*, *Aplnr*, *Ddit4l*) and ECM metalloproteases (i.e. *Adamts8*, *Adamts4*, *Adamts12*) (**Fig. S9d** and **Sup. File 7**). These results are consistent with ORAB inducing metabolic stress and the expression of tissue remodelling mediators in endothelial cells (**Fig. 3h**), and these changes are mostly recovered after DB.

Interestingly, fibroblast cell states show the largest differences across conditions. From the 8 fibroblast cell states, 7 display differential abundance across conditions (ANOVA adj. P-values < 0.05) (**Fig. S10a, b** and **Table S6**). We focused on fibroblast clusters 0, 1, 2 and 3, as each of the other cell states represented < 5% of the total fibroblast nuclei (**Table S6**). Fibroblast clusters 2 and 3 are more abundant in Control than ORAB or DB samples (cluster 2 ANOVA adj. p-value = 1.01e-05, cluster 3 ANOVA adj. p-value = 1.01e-05). Fibroblast cluster 2 was characterised by the expression of genes involved in fatty acid oxidative metabolism (i.e. *Ppara*, *Abcd2*, *Acss3*) and ECM homeostasis (i.e. *Htra1*, *Htra3*, *Fbln5*) (**Fig. S10c** and **Sup. File 8**). The marker genes of fibroblast cluster 3 also suggested that they play a role in ECM homeostasis, with a potential focus in hyaluronan/glycosaminoglycan synthesis (i.e. *Has2*, *Ugdh*, *Gfpt2*) and the assembly of elastic fibers (i.e. *Fbn1*, *Fbln2*, *Emilin2*) (**Fig. S10c** and **Sup. File 8**). Fibroblast cluster 1 is more abundant in ORAB than in Control or DB conditions (ANOVA adj. p-value = 2.23e-07), although the proportion of these cells in DB samples was higher than in Controls (Tukey HSD adj. p-value for DB vs Control = 7.21e-04). Fibroblast cluster 1 is characterised by high expression of genes suggesting increased TGFb signaling (i.e. *Tgfb1*, *Tgfb3*, *Tgfbi*) and collagen secretion (i.e. *Col1a1*, *Col1a2*, *Col3a1*), therefore likely representing a population of activated fibroblasts (i.e. *Postn*, *Acta2*, *Lum*) (**Fig. S10d**). This cluster resembled the TGFb-responsive FB4 cluster described by Litvinukova *et al.*^31^ (**Fig. S10e**). Lastly, the fibroblast cell state analysis identified one fibroblast cluster (cluster 0) enriched in the DB samples compared to all others (ANOVA adj. p-value = 1.28e-04) (**Fig. S10b**). These fibroblasts seem to have recovered the fatty acid metabolism characteristic of controls (i.e. *Abcd2*, *Pdk4*, *Acaa2*), but appear to be executing higher autophagy activity (i.e. *Foxo3*, *Dapk1*, *Ppm1d*) (**Fig. S10c** and **Sup. File 8**).

Overall, the cell state analysis suggested that reverse remodelling triggered by DB might be able to revert CM and endothelial cells to a close to normal state, as only ORAB-specific cell states could be identified for these populations. In contrast, fibroblasts did not fully revert to baseline conditions after DB, and a population specific to reverse remodelling might be retained for at least 2 weeks after DB. This could explain why fibroblasts show the largest number of DEGs and may be the main cell type contributing to the damage-to-recovery gradient.

### 2.5 Identification of ligands driving multi-cellular mechanisms

The apparent reversibility of CM states after DB made us wonder what triggers this reversibility, and particularly whether other cell types influence CMs to achieve this reversibility. To answer this question, we used LIANA+^33^ to predict how cell-cell communication (CCC) networks differed as a result of adverse remodelling (ORAB vs Control) and reverse remodelling (DB vs ORAB). We identified 1,152 and 1,282 deregulated CCC interactions in the adverse remodeling (ORAB vs Control, Sup. File 10) and reverse remodelling (DB vs ORAB, Sup. File 11) comparisons, respectively (**Fig. S11a**). In both cases, over 84% of the identified CCC interactions were mediated by genes encoding secreted ligands (**Fig. S11a**), and fibroblasts were the most frequent senders, being the sender cell in at least 46% of the deregulated CCC interactions (**Fig. S11b**).

To validate our CCC analysis, we started by exploring the deregulated CCC in adverse remodelling, as this has been previously studied^21^. We observed a prominent upregulation of fibroblast-mediated signaling in ORAB compared to Controls (**Fig. S11c**). Most of the fibroblast-mediated interactions upregulated in ORAB are triggered by different types of collagens. We observed an increase in the scar-forming collagen I genes *Col1a1* and *Col1a2*, the fibrillar collagen III gene *Col3a1*, suggested to be deposited during the first stages of adverse remodelling to then be replaced by stiffer collagen I fibers^34^, and the non-fibrillar collagen VI gene *Col6a2*, which has been shown to promote myofibroblast differentiation^35,36^ (**Fig. S11d**).

### 2.6 Multicellular communication identifies a lymphatic RELN-VLDLR signaling axis associated with reverse remodeling

Next, we focused our analysis in understanding which changes in intercellular communication contribute to the recovery of CM homeostasis. We therefore focused our cell-cell communication analysis on interactions selectively enhanced following de-banding, prioritizing those mediated by secreted ligands (see Methods). This approach identified 23 recovery-associated interactions, 19 of which involved secreted ligands (**Fig. S12a** and **Sup. File 12**). Among signals directed toward CMs, only two non-myocyte populations showed increased predicted communication following de-banding: endothelial cells through *Bmp6* and lymphatic endothelial cells (LECs) through *Reln* (**Fig. S12a**). We prioritized RELN for further investigation because its predicted source was confined to the lymphatic compartment and its expression displayed a striking recovery-associated pattern. LEC *Reln* expression decreased significantly during pressure overload (ORAB vs Control log2FC = -0.62, adj. P-value = 8.59e-04) and rebounded following de-banding (DB vs ORAB log2FC = 0.72, adj. P-value = 6.19e-04), returning toward control levels (DB vs Control log2FC = 0.11, adj. P-value = 0.854) (**Fig. 5a**). Mapping *Reln* across the cardiac single-nucleus landscape further revealed a highly restricted cellular distribution, with expression concentrated within the LEC population (**Fig. 5b** and **Fig. S12c**). These findings identified cardiac LECs as a discrete source of RELN and revealed that this lymphatic signal is dynamically suppressed during pressure overload and re-established upon unloading.

**Figure 5.**
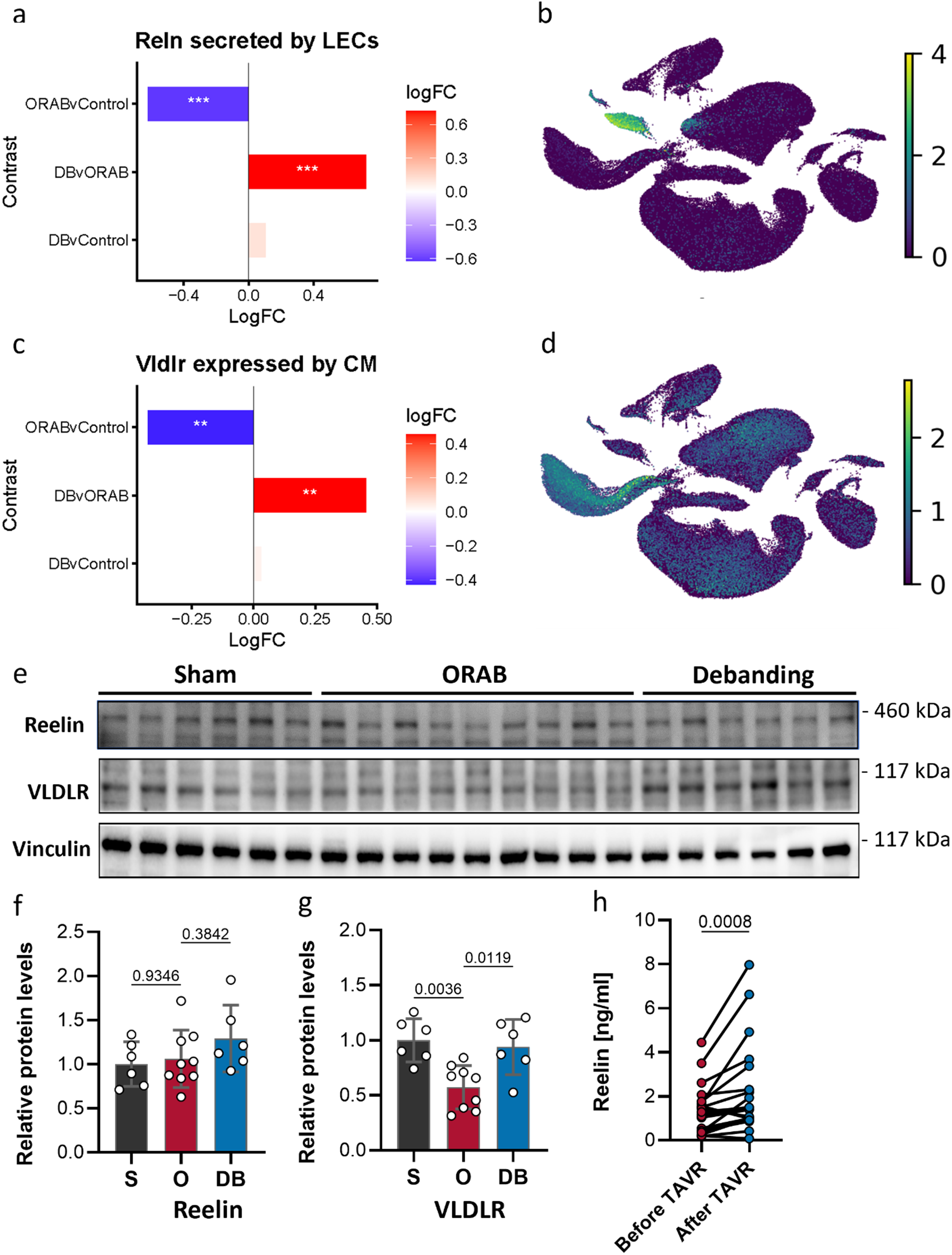
A lymphatic endothelial cell-derived RELN-VLDLR signaling axis is restored during reverse cardiac remodeling. **a)** Differential expression of *Reln* in lymphatic endothelial cells (LECs) across ORAB versus Control, DB versus ORAB, and DB versus Control comparisons derived from single-nucleus RNA sequencing. Bars indicate log fold-change (logFC); asterisks indicate FDR < 0.001 (***). **b)** UMAP visualization of *Reln* expression across cardiac cell populations, demonstrating predominant localization of *Reln* expression to LECs. **c)** Differential expression of *Vldlr* in cardiomyocytes (CMs) across ORAB versus Control, DB versus ORAB, and DB versus Control comparisons. Bars indicate log fold-change (logFC); asterisks indicate FDR < 0.01 (**). **d)** UMAP visualization of *Vldlr* expression across cardiac cell populations, showing prominent expression in cardiomyocytes and additional expression in endothelial cells and fibroblasts. **e)** Representative immunoblots of RELN and VLDLR in cardiac tissue from Control, ORAB, and DB mice; loading control is shown below. **f, g)** Quantification of cardiac RELN (**f**) and VLDLR (**g**) protein abundance in Control, ORAB, and DB mice. n = 6 mice for sham, 9 for ORAB and 6 for de-banding. Data are presented as individual biological replicates as mean ± SEM. Statistical significance was assessed using ordinary one-way ANOVA followed by Tukey’s multiple comparisons test. Exact p-values for the respective comparisons are indicated in the graph. **h)** Circulating RELN levels in patients with severe aortic stenosis before and after transcatheter aortic valve replacement (TAVR), demonstrating significantly increased RELN following relief of pressure overload. Paired samples (n=20) from individual patients are shown. Statistical significance was assessed using Wilcoxon test. Exact p-value is indicated in the graph. Sham, sham-operated control animals (also denoted as ‘S’); ORAB, O-ring aortic banding (also denoted as ‘O’); DB, de-banding; DB, de-banding.

RELN exerts its cellular effects through members of the lipoprotein receptor family, including very-low-density lipoprotein receptor (VLDLR). We therefore examined whether the predicted recipient population displayed corresponding regulation of *Vldlr*. In cardiomyocytes, *Vldlr* expression closely mirrored the behavior of LEC *Reln*: it was significantly reduced following ORAB (ORAB vs Control log2FC = -0.43, adj. P-value = 1.76e-03) and increased after de-banding (DB vs ORAB log2FC = 0.46, adj. P-value = 3.36e-03), again approaching control levels (DB vs Control log2FC = 0.03, adj. P-value = 0.90) (**Fig. 5c**). Across the cardiac cell landscape, *Vldlr* expression was enriched in cardiomyocytes, with additional expression in endothelial cells and fibroblasts (**Fig. 5d** and **Fig. S12c**). Thus, the spatial segregation of ligand and receptor, RELN predominantly in LECs and VLDLR in cardiomyocytes, together with their coordinated regulation across pressure overload and unloading suggested a paracrine LEC-to-cardiomyocyte signaling axis that is re-engaged during reverse remodeling (**Fig. S12d**).

We next examined whether this recovery-associated transcriptional signature was reflected at the protein level. Total myocardial RELN abundance did not differ significantly among Control, ORAB, and DB hearts, although a modest increase was apparent following de-banding (**Fig. 5e,f**). Circulating RELN showed a similar directional pattern, with lower levels following ORAB and an increase after de-banding that did not reach statistical significance (**Fig. S12e,f**). The absence of a significant change in whole-heart RELN protein despite robust regulation of *Reln* within LECs is consistent with the highly restricted cellular origin revealed by single-nucleus profiling and suggests that transcriptional regulation within this relatively small population is not readily captured by bulk myocardial measurements. VLDLR, by contrast, exhibited a pronounced recovery-associated response at the protein level. Myocardial VLDLR abundance was significantly reduced following ORAB and restored after de-banding (**Fig. 5e,g**), closely recapitulating the cardiomyocyte transcriptional pattern. This concordance between cell-type-resolved *Vldlr* expression and myocardial VLDLR protein abundance indicated that unloading restores the receptor component of the predicted RELN signaling axis, potentially re-establishing cardiomyocyte responsiveness to extracellular RELN.

We next asked whether RELN exhibited a corresponding response to therapeutic relief of pressure overload in humans. In patients with severe aortic stenosis undergoing transcatheter aortic TAVR, circulating RELN concentrations increased significantly after the procedure compared with paired pre-TAVR measurements (**Fig. 5h**). Although circulating RELN cannot be assigned to a cardiac or lymphatic source in this setting, its increase following relief of chronic pressure overload provides complementary human evidence that RELN availability is dynamically associated with unloading. Collectively, these findings delineate a recovery-associated RELN-VLDLR signaling spanning the ligand-producing and signal-receiving compartments of the heart. Pressure overload suppresses LEC *Reln* expression together with cardiomyocyte *Vldlr* expression and myocardial VLDLR abundance, whereas unloading reverses these changes. The spatially complementary distribution of RELN and VLDLR, coupled with their coordinated recovery and the increase in circulating RELN following TAVR, raised a key mechanistic question: does re-establishment of this ligand-receptor axis translate into activation of intracellular RELN signaling in the recovering myocardium?

### 2.7 RELN–VLDLR–DAB1 signaling is associated with reverse remodeling and attenuates pathological cardiomyocyte hypertrophy

To explore whether unloading may reinstate a broader signaling program rather than simply restore expression of an individual ligand–receptor pair, we investigated our transcriptomic datasets for genes predicted to be associated with RELN–VLDLR signaling and examined their expression across Control, ORAB, and DB hearts. This analysis revealed a set of 30 genes whose expression in CM was predicted to be regulated by RELN (**Fig. S13a**). We next sought independent evidence for this transcriptional response. Nine candidates spanning stress, growth, inflammatory, and metabolic programs, *Ankrd1, Gdf15, Nfkb1, Bax, Pfkl, Ppargc1a, Cited2, Ccnd2,* and *Etfa*, were selected for validation by qPCR in cardiac tissue (**Fig. S13b–j**). Their expression patterns largely recapitulated those identified by transcriptomic profiling. Genes induced during pressure overload, including *Ankrd1, Gdf15, Nfkb1, Bax,* and *Ccnd1*, downregulated following de-banding, whereas candidates suppressed during ORAB, including *Cited2, Etfa,* and *Ppargc1a*, showed varying degrees of recovery with unloading (**Fig. S13b–j**).

We then asked whether this transcriptional response coincided with biochemical activation of canonical RELN signaling. Engagement of VLDLR by extracellular RELN promotes phosphorylation of the intracellular adaptor DAB1, providing a proximal molecular readout of pathway activation. Immunoblotting of cardiac tissue revealed marked remodeling-dependent regulation of this signaling node (**Fig. 6a** and **Fig. S13k**). DAB1 abundance was reduced following ORAB and remained altered after de-banding, whereas pDAB1 was strongly increased during the recovery phase (**Fig. 6a** and **Fig. S13l**). Notably, pDAB1 abundance in de-banded hearts exceeded that observed in Control hearts, suggesting that unloading does not merely return this pathway to its basal state but is accompanied by pronounced activation of DAB1 signaling. To account for changes in total DAB1 abundance, we next quantified pDAB1 relative to total DAB1. The pDAB1/DAB1 ratio was markedly elevated following de-banding (**Fig. 6a,b**), providing evidence for enhanced DAB1 activation during reverse remodeling and extending the predicted RELN–VLDLR interaction beyond coordinated ligand–receptor expression to activation of its canonical intracellular signaling machinery.

**Figure 6.**
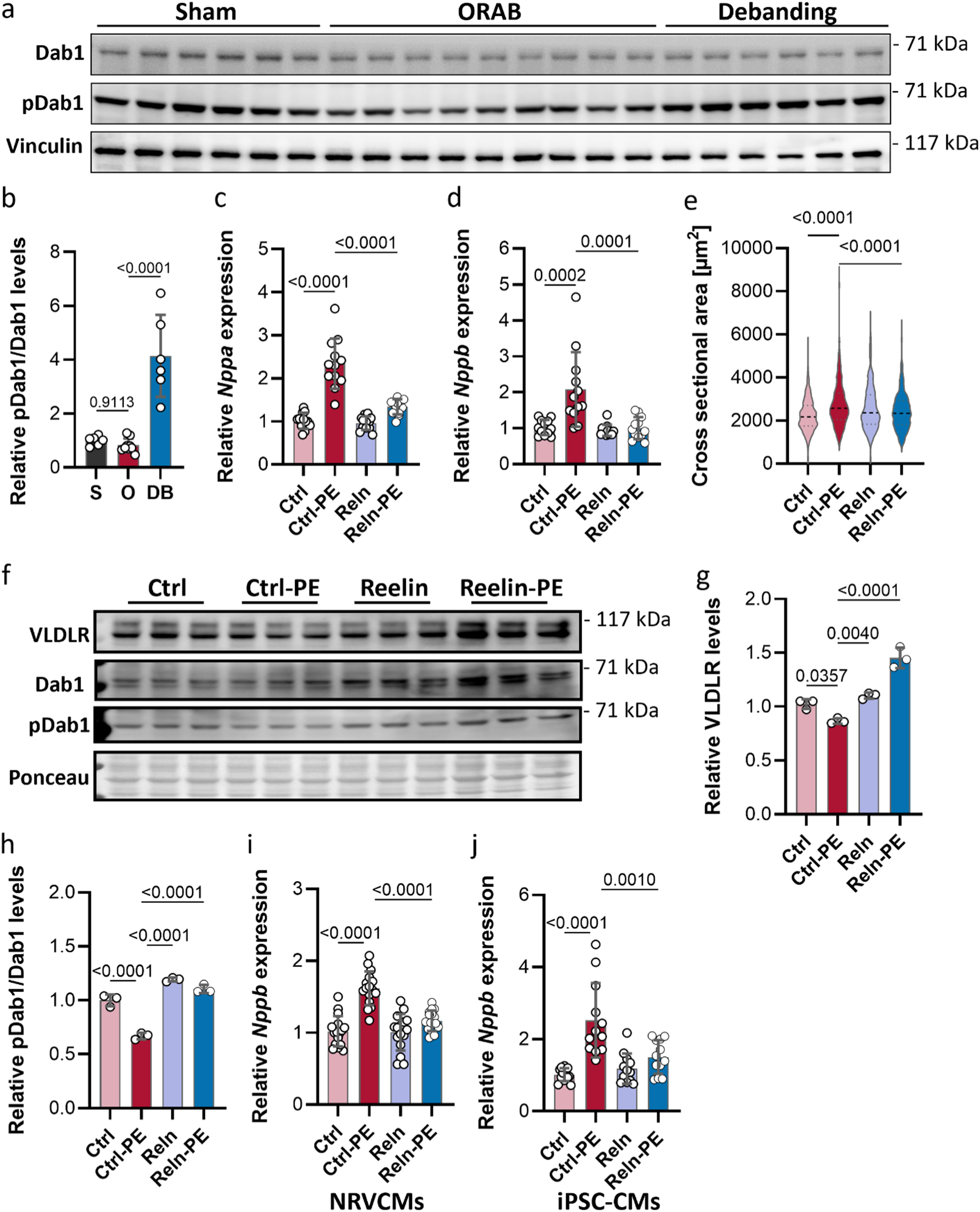
RELN–VLDLR–DAB1 signaling is associated with cardiac recovery and attenuates pathological cardiomyocyte hypertrophy. **a)** Representative immunoblots of phosphorylated DAB1 (pDAB1), total DAB1 and the corresponding loading control in myocardial tissue from Control, ORAB and DB mice. **b)** Densitometric quantification of the pDAB1/DAB1 ratio, showing increased relative DAB1 phosphorylation following de-banding. n = 6 mice for sham, 9 for ORAB and 6 for de-banding. Data are presented as individual biological replicates as mean ± SEM. Statistical significance was assessed using ordinary one-way ANOVA followed by Tukey’s multiple comparisons test. Exact p-values for the respective comparisons are indicated in the graph. **c,d)** Relative expression of the fetal hypertrophic markers *Nppa* (**c**) and *Nppb* (**d**) in neonatal rat ventricular cardiomyocytes (NRVCMs) exposed to conditioned medium derived from empty-vector- or RELN-overexpressing HEK293T cells under basal conditions or following phenylephrine (PE)-induced hypertrophic stress. n = 12 per condition. Data are presented as individual biological replicates as mean ± SEM. Statistical significance was assessed using ordinary one-way ANOVA followed by Tukey’s multiple comparisons test. Exact p-values for the respective comparisons are indicated in the graph. **e)** Quantification of cardiomyocyte surface area in NRVCMs subjected to the indicated conditioned-medium treatments in the presence or absence of PE. Cell surface area was measured in 3 biological replicates with >500 cells per condition. Statistical significance was assessed using ordinary one-way ANOVA followed by Tukey’s multiple comparisons test. Exact p-values for the respective comparisons are indicated in the graph. **f)** Representative immunoblots of VLDLR, total DAB1, pDAB1 and the corresponding loading control in NRVCMs treated with empty-vector- or RELN-enriched conditioned medium under basal and PE-stimulated conditions. **g)** Densitometric quantification of VLDLR abundance in NRVCMs following the indicated treatments. **h**) Quantification of the pDAB1/DAB1 ratio, showing restoration of relative DAB1 phosphorylation following exposure to RELN-enriched conditioned medium under PE-induced hypertrophic stress. For ‘g’ and ‘h’, n = 3 biological replicates per group. Data are presented as individual biological replicates with mean ± SEM. Statistical significance was assessed using ordinary one-way ANOVA followed by Tukey’s multiple comparisons test. Exact p-values for the respective comparisons are indicated in the graph. **i**) Relative *Nppb* expression in NRVCMs treated with a commercially available biologically active RELN fragment under basal conditions or following PE stimulation. **j**) Relative *NPPB* expression in human induced pluripotent stem cell-derived cardiomyocytes (hiPSC-CMs) treated with the biologically active RELN fragment under basal conditions or following PE stimulation. For ‘i’ and ‘j’, n = 12 biological replicates per group. Data are presented as individual biological replicates with mean ± SEM. Statistical significance was assessed using ordinary one-way ANOVA followed by Tukey’s multiple comparisons test. Exact p-values for the respective comparisons are indicated in the graph. Sham, sham-operated control animals (also denoted as ‘S’); ORAB, O-ring aortic banding (also denoted as ‘O’); DB, de-banding; RELN, Reelin; VLDLR, very-low-density lipoprotein receptor; DAB1, Disabled-1; PE, phenylephrine.

The recovery-associated activation of DAB1 observed in vivo suggested that restoration of RELN–VLDLR signaling may directly influence cardiomyocyte state. We therefore tested whether extracellular RELN is sufficient to engage VLDLR–DAB1 signaling in cardiomyocytes and attenuate pathological hypertrophic remodeling. To model exposure to cell-secreted RELN, we overexpressed RELN in HEK293T cells and collected the resulting conditioned medium. Immunoblotting confirmed robust enrichment of RELN in culture supernatant from RELN-overexpressing cells compared with empty-vector controls (**Fig. S13m**). The conditioned media were subsequently applied to neonatal rat ventricular cardiomyocytes (NRVCMs) under basal conditions or during phenylephrine (PE)-induced hypertrophic stress. PE treatment induced a robust hypertrophic transcriptional response, characterized by increased expression of the fetal genes *Nppa* and *Nppb*. Exposure to RELN-enriched conditioned medium significantly attenuated the PE-induced increase in both markers compared with conditioned medium from empty-vector-transfected cells (**Fig. 6c,d**). This effect extended beyond transcriptional markers of stress. Consistent with its hypertrophic response, PE increased cardiomyocyte surface area, whereas RELN-enriched conditioned medium significantly reduced PE-induced cardiomyocyte enlargement (**Fig. 6e**). Importantly, RELN-enriched conditioned medium did not adversely affect cell viability, as assessed by MTT assay (**Fig. S13n**), indicating that the reduction in hypertrophic gene expression and cell size was not secondary to compromised cardiomyocyte survival. Together, these findings demonstrate that exposure to RELN-enriched conditioned medium counteracts both molecular and morphological features of pathological cardiomyocyte hypertrophy.

We next asked whether this phenotypic response was accompanied by engagement of the same signaling pathway that was activated during reverse remodeling in vivo. Immunoblotting revealed increased VLDLR abundance in NRVCMs exposed to RELN-enriched conditioned medium, an effect that was maintained in the presence of PE (**Fig. 6f,g**). RELN exposure was also associated with increased abundance of DAB1 and its phosphorylated form, pDAB1 (**Fig. 6f** and **Fig. S13o,p**). Notably, PE reduced relative DAB1 activation, whereas exposure to RELN-enriched conditioned medium restored the pDAB1/DAB1 ratio despite continued hypertrophic stimulation (**Fig. 6f,h**). Thus, extracellular RELN exposure not only attenuated the hypertrophic phenotype but was also accompanied by increased VLDLR abundance and DAB1 phosphorylation in cardiomyocytes. This response closely aligned with the restoration of VLDLR expression and DAB1 phosphorylation observed in de-banded hearts, linking the recovery-associated signaling signature identified in vivo to a direct cardiomyocyte response to RELN-enriched conditioned medium.

As an orthogonal validation, we tested whether a defined, commercially available biologically active RELN fragment could reproduce the effects observed with cell-secreted RELN. Treatment with the RELN fragment attenuated PE-induced *Nppb* expression in NRVCMs (**Fig. 6i**). This response was conserved in human induced pluripotent stem cell-derived cardiomyocytes (hiPSC-CMs), in which RELN similarly suppressed PE-induced *NPPB* expression, without impacting cell viability (**Fig. 6j** and **Fig. S13q**). The concordant effects of RELN-enriched conditioned medium and a defined active RELN fragment across complementary cardiomyocyte models provide independent support for an antihypertrophic effect of extracellular RELN and extend this observation to a human cellular context.

Collectively, these experiments connect the recovery-associated RELN–VLDLR signaling architecture identified in vivo with the effects of extracellular RELN on cardiomyocytes. Together with the response observed following treatment with a defined RELN fragment in rat and human cardiomyocytes, these findings support a model in which LEC-derived RELN contributes to reverse cardiac remodeling through a VLDLR–DAB1-associated signaling response that favors recovery from pathological hypertrophic stress.

## 3. Discussion

In this study, we characterize reverse cardiac remodeling as a heterogeneous multicellular recovery process rather than a simple return to the non-failing state. By integrating phenotypic characterization with bulk and single-nucleus transcriptomics, we demonstrate that hemodynamic unloading selectively reverses pathological programs while leaving others unresolved. Cardiomyocytes and endothelial cells largely reacquire homeostatic transcriptional features, whereas fibroblasts retain activated states, and inflammatory and mitochondrial abnormalities persist across cardiac cell types. Multicellular factor analysis further distinguishes reversible stress-associated responses from persistent pathological programs, revealing that functional improvement and molecular recovery are not necessarily concordant. Beyond defining these divergent recovery trajectories, our integrated analyses identify candidate intercellular signals that may contribute to myocardial adaptation following unloading, with lymphatic endothelial cell (LEC)-derived RELN providing one example supported by targeted biochemical and functional investigation. Together, these findings establish a framework for understanding the cellular heterogeneity of reverse remodeling and prioritizing recovery-associated pathways for future mechanistic studies.

Our observations support an emerging view that functional recovery in heart failure does not necessarily equate to complete biological normalization. Clinical studies of patients supported by left ventricular assist devices (LVADs) similarly demonstrate improvements in cardiac function and geometry despite persistent fibrosis, metabolic abnormalities, and molecular features of heart failure^37–39^. More recent work has shown that recovered myocardium remains transcriptionally distinct from non-failing myocardium and can retain mitochondrial and epigenetic abnormalities after unloading^19,40,41^. This distinction is clinically relevant because relapse can occur even after apparent recovery. Long-term follow-up of TRED-HF demonstrated that recurrent dysfunction continued to accumulate after withdrawal of therapy in many patients^40,42^. Although the experimental and clinical settings differ, these observations collectively suggest that reverse remodeling may represent a state of functional improvement accompanied by residual molecular vulnerability rather than complete restoration of myocardial homeostasis.

### Cell-type-specific recovery reveals persistent stromal and inflammatory abnormalities

In this study, single-nucleus RNA sequencing revealed marked differences in the extent of recovery across cardiac cell types. Cardiomyocytes and endothelial cells approached control-like transcriptional states after de-banding, whereas fibroblasts retained activated and remodeling-associated programs. This divergence provides a potential explanation for why tissue-level recovery remains incomplete despite substantial normalization of cardiomyocyte transcriptional states. The recovery of endothelial programs is particularly relevant in light of recent evidence that mechanical unloading can reverse microvascular senescence and remodel the cardiac vascular niche, including endothelial, pericyte, and fibroblast populations^43^. Together, these observations suggest that unloading promotes recovery of important vascular and myocardial cell states while leaving distinct stromal abnormalities unresolved.

Persistent fibroblast activation has also been described in human heart failure and following myocardial injury^20,39,44–47^. Recent studies further indicate that fibroblasts can impair cardiomyocyte function through paracrine signaling independently of bulk fibrosis, suggesting that fibroblast activation states may be more informative than collagen abundance alone^48^. This is consistent with our observation that fibroblast abnormalities persisted despite partial regression of histological fibrosis. Notably, the fibroblast population enriched after de-banding may represent a distinct remodeling-associated state rather than simple persistence of the pressure-overload phenotype. These findings emphasize that regression of extracellular matrix deposition and normalization of stromal cell states are not necessarily equivalent processes. Persistent fibroblast programs may therefore represent an important component of the residual myocardial pathology observed after unloading, although their direct contribution to limiting recovery remains to be established.

The multicellular analyses extended these observations beyond individual cell types by distinguishing transcriptional programs with divergent responses to unloading. While a metabolic-stress-associated program showed substantial recovery, another program characterized by interferon signaling, inflammation, and altered mitochondrial metabolism remained dysregulated. The association between inflammatory and metabolic abnormalities is consistent with mechanistic studies demonstrating that inflammatory signaling can reprogram cardiomyocyte metabolism and suppress oxidative pathways, including through IFN- and IRF3-dependent inhibition of PGC-1α^49,50^. These findings provide biological context for the persistence of inflammatory and mitochondrial abnormalities within the same multicellular program. Notably, inflammatory signatures remained detectable despite reduced myeloid abundance, suggesting that inflammatory transcriptional activity after unloading is not explained solely by immune-cell expansion and may also involve resident cardiac populations. Such persistent programs could reflect a form of molecular memory following chronic biomechanical stress. However, whether they actively restrict subsequent recovery or gradually resolve with longer unloading remains an important question for future investigation.

### Intercellular communication identifies candidate recovery-associated pathways

Beyond defining the cellular and transcriptional architecture of reverse remodeling, our integrated analyses identified candidate intercellular interactions associated with myocardial recovery. Among these, we investigated lymphatic endothelial cell (LEC)-derived RELN and cardiomyocyte VLDLR as an example of how transcriptomic inference can guide functional evaluation of recovery-associated pathways. Both *Reln* and *Vldlr* were suppressed during pressure overload and restored following de-banding, accompanied by recovery of myocardial VLDLR protein. This coordinated response suggests that unloading may restore a lymphatic–cardiomyocyte signaling interaction within the broader multicellular recovery program.

RELN is a secreted glycoprotein that signals through VLDLR and LRP8 (ApoER2) to promote phosphorylation of the intracellular adaptor DAB1^51,52^. Beyond its established neurological functions, RELN has been implicated in tissue organization, vascular regulation, and immune responses^53,54^. Cardiac lymphatics are increasingly recognized as regulators of myocardial homeostasis and repair^24–26^, with previous studies identifying lymphatic-derived RELN as a mediator of cardiomyocyte survival and repair after injury and implicating VLDLR-dependent signaling in cardiac growth^24,55^. RELN–VLDLR signaling has also been linked to YAP-dependent cardiomyocyte responses^24,55^, consistent with broader roles of YAP in cardiac growth and stress adaptation^56,57^. Together, these findings provided us with biological context for investigating RELN during recovery from chronic pressure overload.

Consistent with the predicted interaction, de-banding was accompanied by increased myocardial DAB1 phosphorylation relative to total DAB1, suggesting enhanced activation of a canonical RELN-associated signaling node. Functionally, RELN-enriched conditioned medium attenuated hypertrophic gene expression and cardiomyocyte enlargement while increasing VLDLR abundance and DAB1 phosphorylation. A defined biologically active RELN fragment similarly suppressed hypertrophic gene induction in rat and human cardiomyocytes, providing complementary support for an antihypertrophic response to extracellular RELN. In parallel, circulating RELN increased in patients following TAVR, extending the recovery-associated observations to a clinical unloading setting. However, the origin of circulating RELN remains unresolved, particularly given its reported expression also in calcific aortic valve tissue^58^.

Together, these findings illustrate how multicellular transcriptomic analyses can identify candidate recovery-associated signals with experimentally demonstrable effects on cardiomyocyte remodeling. Although the coordinated RELN–VLDLR response, DAB1 phosphorylation, and functional observations support the biological relevance of this pathway, its causal contribution to reverse remodeling in vivo remains to be established. RELN therefore represents one promising example within a broader network of recovery-associated intercellular interactions, rather than a singular determinant of myocardial recovery.

### Limitations

Several limitations should be acknowledged. First, the ORAB/de-banding model captures a defined etiology and recovery interval in male mice, limiting generalizability and the ability to distinguish persistent from permanently irreversible remodeling programs. Second, multicellular and cell–cell communication analyses identify associations rather than causal relationships. Although the RELN– VLDLR–DAB1 pathway was supported by biochemical and functional experiments, its receptor dependence and contribution to reverse remodeling in vivo require further investigation. Finally, the tissue origin of circulating RELN following TAVR remains unresolved.

Despite these limitations, our study establishes reverse cardiac remodeling as a coordinated yet incomplete multicellular recovery process, characterized by divergent cell-type-specific responses and persistent pathological programs. By identifying candidate recovery-associated intercellular signals, exemplified by the functionally investigated RELN–VLDLR–DAB1 pathway, our findings provide a framework for future mechanistic studies. Ultimately, achieving more complete and durable myocardial recovery may require both resolution of persistent pathological programs and reinforcement of endogenous pro-recovery signaling.

## 4. Methods

### 4.1 Animal model, surgery, functional phenotyping and histology

#### Ethics

All animal experiments were carried out stringently following international and institutional ethical guidelines, replicated in two independent cohorts, single-blinded and approved by the Regierungspräsidium Karlsruhe Institutional Animal Care and Use Committee (G-174/23, Baden-Württemburg, Germany).

#### Animal model

All mice were purchased from Janvier Labs (France) with a C57BL/6N wild-type background (N=36 and N=15 first and second cohort, respectively). The mice were housed at the Klinisch Experimenteller Bereich mouse facility of Heidelberg University, randomized after arrival and single-housed on a 12:12-hour light-dark cycle from 06:00 to 18:00 at 22±1 °C as well as constant humidity of 50-60% with ad libitum access to food (Altromin Rod 16 or Rod 18) and water.

#### ORAB and de-banding mouse model

O-ring aortic banding (ORAB) was performed by placing a nitrile o-ring around the transverse aorta using a previously described technique^59^. Nitrile o-rings with an inner diameter of 0.46 mm (Apple Rubber, Lancaster, NY, USA) were used to induce pressure overload in 12-week-old male C57BL/6N mice. Mice that underwent ORAB were anesthetized with isoflurane, intubated, and ventilated with 1.5% isoflurane in oxygen. A 1 cm longitudinal incision was made in the skin of the upper left thorax; the pectoralis muscles were retracted, the intercostal muscles bluntly dissected, and the ribs gently spread. After retracting the left lung and thymus, the transverse aorta was carefully isolated using micro forceps. The open o-ring was positioned around the vessel and closed by tying the attached sutures. Excess suture material was removed, the pectoral muscles were repositioned to cover the incision, and the skin was closed with continuous sutures. Buprenorphine (0.1 mg/kg, three times daily, ten injections total) was administered for intra- and postoperative analgesia. For de-banding, mice were anesthetized and the transverse aorta was accessed following the same approach as for the banding procedure. The sutures closing the o-ring were cut, the o-ring was removed, the wound was closed and analgesia was provided as described above. Sham-operated animals underwent the same procedure without placement (Sham ORAB) or removal (Sham De-banding) of an o-ring.

#### Cardiac echocardiography

Echocardiography was performed in non-anesthetized mice using a Visual Sonics Vevo 2100 setup equipped with a MX550D transducer. B- and M-mode images were obtained from left ventricular parasternal long views and short-axis views at the mid-papillary muscle level. Analysis was performed using Visual Sonics Vevo LAB software’s LV trace tool with the Teicholz equation, based on at least five contraction cycles.

#### Histology

After organ harvest, mouse hearts were put in 4% PFA for 48h and then transferred into 70% ethanol. The hearts were embedded in paraffin using the HistoCore PEARL Tissue Processor and Arcadia Embedding Center (both from Leica Biosystems, Germany). Sections of 5 µm thickness were cut, mounted on slides, dried on a hotplate (30 min), followed by overnight drying in an incubator (37°C). Sections were dewaxed for 10 mins twice using ROTI®Histol (Carl Roth GmbH + Co. KG, Germany), rehydrated with a descending alcohol series (100% for 2 minutes, 96% for 2 minutes, and 70% for 2 minutes) and in the end placed in distilled water.

H&E staining was performed using H&E fast staining kit (Carl Roth GmbH + Co. KG, Germany) in accordance with the manufacturer’s protocol. Sirius Red staining was performed by immersing slides in Picro-Sirius Red Solution (ScyTek Laboratories, USA) for 30 minutes with subsequent differentiation in acidified water (1% acetic acid, two times). Dehydration with an ascending alcohol series (70% for 2 minutes, 96% for 2 minutes, and 100% for 2 minutes) was performed, cleared in ROTI®Histol and mounted with 35 µl EUKITT® neo mounting medium (ORSAtec GmbH, Germany). Images of the sections were taken using ZEISS Axioscan slide scanner (ZEISS, Germany).

H&E (cross sectional area) and Sirius Red (fibrotic area) analysis was done in Fiji (ImageJ 1.54f, USA, Java 1.8.0.322 (64-bit)). In brief, snapshots of H&E full resolution images were taken in similar areas for each of the samples (three 10x and seven to eight 20x magnified snapshots) with the ZEISS ZEN lite software (ZEISS, Germany). A cross-sectional area of 15-20 cells per image was measured in Fiji using the “set scale” function and the “freehand selection” tool. For fibrosis analysis, the ROI manager was opened and a defined snapshot form was generated using the “selection” and “specify” function (5000 width & length, oval ROI). This form was placed over the Sirius Red sections in similar areas for each sample, 11 times in total, always connected to the last form, so that the placed 11 forms covered most of the heart section as possible. Each of those snapshots was analyzed single-blinded for the amount of fibrotic area compared to non-fibrotic area in percent, determined by an image specific threshold that defined the background.

### 4.2 Isolation and culture of neonatal rat cardiomyocytes

Neonatal rat ventricular cardiomyocytes (NRVCMs) were isolated from 1- to 2-day-old Sprague-Dawley rat pups using a modified protocol adapted from Miltenyi Biotec’s Neonatal Heart Dissociation Kit (mouse/rat, cat#130-098-373) and Neonatal Cardiomyocyte Isolation Kit (rat, cat#130-105-420). Briefly, neonatal hearts were dissected in ice-cold ADS buffer (197.2 mM HEPES, 1163.6 mM NaCl, 55.5 mM glucose, 94.2 mM NaH₂PO₄, 8.3 mM MgSO₄, 53.6 mM KCl, pH 7.4). Connective tissue was removed, hearts were minced and enzymatically dissociated in a gentleMACS Octo Dissociator (37 °C, 56 min) using a two-step enzyme mixture (enzyme Mix I contains Enzyme P and Buffer X; enzyme Mix II contains Enzyme A, Enzyme D, and Buffer Y). Filtering of the cell suspension through a 70-µm Cellstrainer was followed by centrifugation and red blood cell depletion using Anti-Red Blood Cell Microbeads. Cardiomyocytes were purified via magnetic-activated cell sorting (MACS) columns, resuspended in DMEM/F12 medium supplemented with 10% fetal calf serum (FCS) and 1% penicillin-streptomycin-glutamine, and counted using trypan blue exclusion. Cells were seeded in DMEM containing 10% fetal calf serum (FCS), 2 mM penicillin/streptomycin, and L-glutamine (PAA Laboratories, Austria) and all experiments were conducted 24h post seeding.

### 4.3 Differentiation of human induced pluripotent stem cells into cardiomyocytes

Human induced pluripotent stem cell-derived cardiomyocytes (hiPSC-CMs) were obtained cryopreserved from the StemCell Hub (Dr. Timon Seeger). For differentiation, Matrigel-coated 6-well plates were prepared 2h before seeding. Cells were thawed on ice, gently centrifuged at 300 × g for 3 min, and resuspended in hE8 medium with ROCK inhibitor (Y-27632 Tocris, England) before plating. Cultures were maintained at 37 °C in a humidified incubator with 5% CO_2_. Differentiation of cardiomyocytes was carried out using a defined protocol^60^. Cells were treated with 3 ml CDM3 medium containing CHIR99021 (CHIR, MedChemExpress, USA) on day 0. On day 2, medium was changed to CDM3 alone. CDM3 was supplemented with IWP2 (Tocris, England) on days 3 and 4 and changed back to CDM3 alone from days 5 to 6. Starting day 7, cells were cultured in B27 supplemented with insulin (B27 + ins, Gibco, England) followed by metabolic selection by glucose-depleted CDM3 on days 10 to 12. Cells were maintained in B27+ and passaged on day 15. All experiments were conducted using hiPSC-CMs at 40 days post-differentiation.

### 4.4 RNA isolation and quantitative RT-PCR

Total RNA from mouse hearts, NRVCMs and hIPSCs samples for qPCR was extracted with TRIzol™ reagent (Thermo Fisher Scientific, USA, cat# 15596026) and one microgram of DNA-free RNA was reverse-transcribed using the LunaScript™ RT SuperMix Kit (New England Biolabs, cat#E3010), both according to the manufacturer’s protocol. PowerTrack™ SYBR Green Mastermix (Applied Biosystems, USA, cat# A46111) was used for qRT-PCR and performed on a Roche LightCycler 480. The qRT-PCR protocol included an initial denaturation step at 95 °C for 10 min followed by 40 amplification cycles of 15 seconds at 95 °C and 60 seconds at 60 °C. Rpl32 as a housekeeping gene served as internal control and the ΔΔCt method was used for relative quantification. NRVCMs and hIPSCs experiments were repeated thrice and conducted in 4 or 6 replicates. All primers that were used are summarized in **Sup. File 12**.

### 4.5 In vitro validation of ligand receptor candidates

Following recombinant proteins were used for the in vitro validation: Recombinant Human Reelin (R&D Systems, USA, cat#8546-MR). Hypertrophy was induced by phenylephrine treatment (PE, 50 µM, Sigma-Aldrich, cat# P6126-10G). NRVCMs were seeded at 500k/per well in a 12-well plate, while hiPSC-CMS were seeded at 250k/per well in a 24-well plate. NRVCMs were serum starved for 48 h before continuing with the treatments. hiPSCs-CMs were taken from the StemCell Hub after media change and experiments were conducted 24h later. Except for the respective media (serum free DMEM with 2 mM penicillin/streptomycin & L-glutamine for NRVCMs and B27+ for hiPSC), the experimental setup was the same for both in vitro validation approaches: Cells were washed with PBS and treated with PE, recombinant protein or PE plus recombinant protein for 24h. Vehicle treated cells were used as negative control.

#### Plasmid preparation and generation of RELN-enriched conditioned medium

The full-length, untagged mouse *Reln* expression plasmid pCrl was obtained from Addgene (plasmid #122443; RRID), originally deposited by Tom Curran^61^. Plasmid-containing *Escherichia coli* DH5α were streaked onto LB agar supplemented with ampicillin (100 µg/mL) and incubated overnight at 37 °C. A single colony was expanded in liquid LB medium containing ampicillin under continuous shaking at 37 °C. Plasmid DNA was isolated using the QIAGEN Plasmid Maxi Kit (QIAGEN, Netherlands, cat#12165) according to the manufacturer’s instructions. DNA concentration and purity were assessed using a DeNovix SD-11 FX instrument.

HEK293T cells were maintained in Advanced DMEM (Thermo Fisher Scientific) supplemented with 4% fetal calf serum (FCS), 4.5 g/L glucose, 110 mg/L sodium pyruvate, 100 µg/mL penicillin, 100 µg/mL streptomycin and 2 mM L-glutamine. Cells were seeded at a density of 2 × 10⁶ cells per 10-cm dish and transfected at approximately 70–80% confluence using Lipofectamine 2000 (Thermo Fisher Scientific). Transfection was performed in 10 mL antibiotic-free DMEM supplemented with 10% FCS, using 1.16 µg/mL plasmid DNA and 2.9 µL/mL Lipofectamine 2000. Plasmid DNA and transfection reagent were separately incubated for 5 min at room temperature, combined for an additional 20 min and added dropwise to the cells. After 6 h, the transfection medium was replaced with phenol red-free, serum-free and antibiotic-free HyClone DMEM. Conditioned medium was collected 48 h after transfection and subsequently used for cardiomyocyte treatment. RELN enrichment was confirmed by immunoblotting.

### 4.6 In vitro validation of RELN signaling

To investigate the effects of extracellular RELN on cardiomyocyte hypertrophy, neonatal rat ventricular cardiomyocytes (NRVCMs) and human induced pluripotent stem cell-derived cardiomyocytes (hiPSC-CMs) were treated with RELN-enriched conditioned medium or recombinant human RELN (R&D Systems, USA, cat#8546-MR). Recombinant RELN was applied at a concentration of 150 ng/mL, whereas conditioned medium was diluted 1:1 with the respective culture medium. Hypertrophic stress was induced using phenylephrine (PE; 50 µM; Sigma-Aldrich, cat#P6126-10G).

NRVCMs were seeded at a density of 2 × 10⁵ cells per well in 24-well plates and serum-starved for 48 h before treatment. hiPSC-CMs were seeded at 2.5 × 10⁵ cells per well in 24-well plates and treated 24 h after medium replacement. Experiments were performed in serum-free DMEM supplemented with penicillin–streptomycin and L-glutamine for NRVCMs, or B27-supplemented medium for hiPSC-CMs. Cells were washed with phosphate-buffered saline (PBS) and treated with PE, RELN-containing preparations or their combination for 24 h. Corresponding control treatments were included for each experimental condition. Following treatment, cells were harvested for RNA or protein analysis, or processed for immunofluorescence staining and viability measurements as described below.

### 4.7 Cardiomyocyte viability assay

NRVCM viability was assessed using an MTT-based colorimetric assay (Cell Proliferation Kit I, Roche Applied Science, Germany). Cells were cultured in 24-well plates and subjected to the same experimental conditions as those used for the conditioned-medium experiments. MTT labeling reagent was added at 10% of the culture medium volume, followed by incubation for 4 h at 37 °C. Subsequently, solubilization solution was added at ten times the volume of the MTT labeling reagent, and plates were incubated overnight at 37 °C to ensure complete dissolution of the formazan crystals.

An aliquot of 200 µL from each well was transferred to a 96-well plate, and absorbance was measured using an Infinite M200 PRO microplate reader (Tecan, Switzerland). Cell viability was expressed as a percentage of the corresponding control condition. Experiments were performed using six replicates per condition and repeated in three independent experiments.

### 4.8 Immunofluorescence staining and cardiomyocyte size analysis

For assessment of cardiomyocyte size, NRVCMs were seeded at a density of 2 × 10⁵ cells per chamber in collagen-coated µ-Slide 4 Well chambers (ibidi, Germany). Following treatment, cells were washed three times with PBS and fixed with 4% paraformaldehyde for 30 min at room temperature. After washing, cells were blocked and permeabilized in PBS containing 2.5% bovine serum albumin and 0.1% Triton X-100 for 60 min at room temperature.

Cells were incubated overnight at 4 °C with a mouse monoclonal anti-α-actinin antibody (1:200; Sigma-Aldrich), followed by an Alexa Fluor 488-conjugated secondary antibody (1:400; Thermo Fisher Scientific) for 90 min at room temperature. Subsequently, cells were stained with phalloidin (1:400; Thermo Fisher Scientific) for 60 min and DAPI (1:1,000) for 8 min at room temperature, with four PBS washes between staining steps.

Fluorescence images were acquired using a BZ-X1000 fluorescence microscope equipped with a 20× objective (Keyence, Japan). Cardiomyocyte planar surface area was quantified using Fiji (ImageJ). Cell boundaries were identified by intensity-based thresholding, and individual mononucleated cardiomyocytes with clearly distinguishable borders were selected using the Wand tool. The area of each resulting region of interest was measured and used for quantitative analysis.

### 4.9 Protein extraction and immunoblotting

Protein lysates from mouse left ventricular tissue were prepared by mechanical homogenization using a Precellys 24 Touch Homogenizer (Bertin Technologies, France) with 2.8-mm and 1.4-mm zirconium oxide beads. Tissue was homogenized in RIPA buffer containing 50 mM Tris, 150 mM NaCl, 1% Nonidet P-40, 0.5% sodium deoxycholate and 1% SDS, supplemented with protease inhibitors (4%) and phosphatase inhibitors II and III (1%; Roche Applied Science). Homogenization was performed for two cycles of 30 s at 6,500 rpm.

NRVCM and HEK293T cell lysates were prepared using RIPA buffer and subjected to three or ten freeze– thaw cycles, respectively. Tissue and cell lysates were clarified by centrifugation at 12,000 × g for 20 min at 4 °C. Protein concentrations were determined using the DC Protein Assay (Bio-Rad, Germany) with bovine serum albumin as a standard.

Protein samples were separated by SDS–PAGE. RELN was resolved using a gel system suitable for high-molecular-weight proteins, whereas other proteins were separated using 4–20% Novex Tris-Glycine Plus Midi Protein Gels (Thermo Fisher Scientific). Proteins were transferred to nitrocellulose membranes and incubated overnight with primary antibodies against RELN (1:500; Novus Biologicals, NB600-1081), phosphorylated DAB1 (pDAB1; 1:1,000; Thermo Fisher Scientific, BS-3115R), total DAB1 (1:500; Novus Biologicals, NBP2-92015), VLDLR (1:1,000; Novus Biologicals, NBP1-78162) and vinculin (1:1,000; Cell Signaling Technology, #4650).

Membranes were incubated with horseradish peroxidase-conjugated anti-mouse or anti-rabbit secondary antibodies (1:10,000; Biozol, cat#715-035-150 and 711-035-152, respectively). Protein bands were visualized using Amersham ECL Select chemiluminescence reagent (Cytiva) and imaged using a ChemiDoc MP Imaging System (Bio-Rad). Densitometric analysis was performed using Fiji. Protein abundance was normalized to vinculin or total lane protein, as appropriate for the respective experiment. Relative DAB1 activation was calculated as the pDAB1/DAB1 ratio.

### 4.10 Human plasma samples and RELN quantification

Human plasma samples were obtained from 20 patients with severe aortic stenosis undergoing TAVR. Samples had been collected as part of an independent clinical study entitled “Kardiale Amyloidose bei Patienten mit hochgradiger Aortenklappenstenose und transfemoralem Aortenklappenersatz (TAVR)”, approved by the Ethics Committee of the Medical Faculty of Heidelberg University (S-587/2019). All participants provided written informed consent.

Peripheral blood was collected into lithium-heparin tubes (S-Monovette Lithium Heparin Gel LH, 7.5 mL; Sarstedt, Germany). Plasma was separated by centrifugation at 2,500 rpm for 10 min at room temperature and stored at −80 °C until analysis. Plasma aliquots were thawed and diluted 1:10 before measurement of RELN concentrations using a commercially available human RELN ELISA kit (Novus Biologicals, cat#NBP2-82529), according to the manufacturer’s instructions. Absorbance was measured at 450 nm using an Infinite M200 PRO microplate reader (Tecan, Switzerland).

### 4.11 Bulk RNA-seq RNA extraction, library preparation and sequencing

Total RNA for bulk RNA sequencing was isolated using the RNeasy Plus Universal Mini Kit (Quiagen, Netherlands, cat#73404) following the manufacturer’s instructions. RNA from each sample was quantified using a DeNovix SD-11 FX instrument and sent to Novogene for RNA seq. Briefly, total amounts and integrity of RNA were assessed using the RNA Nano 6000 Assay Kit of the Bioanalyzer 2100 system (Agilent Technologies, USA). A total of 1 μg RNA per sample was used as input material for the mRNA library preparation. Strand-specific libraries were generated using NEBNext® UltraTM RNA Library Prep Kit for Illumina® (NEB, USA) following manufacturer’s recommendations and index codes were added to attribute sequences to each sample. Ribosomal RNA was depleted from total RNA. Fragmentation was carried out using divalent cations under elevated temperature in NEBNext First Strand Synthesis Reaction Buffer(5X). First strand cDNA was synthesized using random hexamer primer and M-MuLV Reverse Transcriptase (RNase H-). Second strand cDNA synthesis was subsequently performed using DNA Polymerase I and RNase H with the incorporation of dUTP instead of dTTP. Remaining overhangs were converted into blunt ends via exonuclease/polymerase activities. After adenylation of 3’ ends of DNA fragments, NEBNext Adaptors with hairpin loop structure were ligated for hybridization. In order to select cDNA fragments of preferentially 250-300 bp in length, the library fragments were purified with AMPure XP system (Beckman Coulter, USA). USER Enzyme (NEB, USA) was used to digest the second strand with size-selected, adaptor-ligated cDNA at 37°C for 15 min followed by 5 min at 95 °C before PCR. Subsequent PCR was performed with Phusion High-Fidelity DNA polymerase, Universal PCR primers and Index (X) Primer. The resulting library was quantified by Qubit2.0 Fluorometer, then diluted to 1.5ng/ul, and the insert size of the library is detected by Agilent 2100 bioanalyzer. qRT-PCR was used to accurately quantify the effective concentration of the library (> 2nM).

The libraries were pooled based on effective concentration to ensure equal data output and sequenced on the Illumina NovaSeq 6000 platform using a paired-end 150 bp (2×150bp) strategy. Raw data in FASTQ format were processed using custom scripts to remove adapter sequences, poly-N and low-quality reads. Quality metrics, including Q20, Q30, and GC content, were calculated for the resulting clean data. All subsequent analyses were performed using these high-quality filtered reads. The genome index was constructed and paired-end clean reads were aligned to the reference genome (ensembl_103_mus_musculus_grcm39_toplevel) using HISAT2 (v2.0.5). Gene expression levels were quantified using StringTie (v1.3.3b).

### 4.12 Bulk RNA-seq analysis

Raw read counts were filtered to remove genes with less than 10 counts in less than 3 samples. Gene expression was normalised using the blind variance stabilizing transformation (VST) before visualisation and calculation of the euclidean distance between samples. Euclidean distance between samples was calculated using the R function dist(). Euclidean distances between groups were compared using the Kruskal-Wallis test, followed by a pairwise Wilcox test and Bonferroni adjustment of p-values. PCA was calculated on the 500 most variable genes. For further analyses, the two control groups (Sham-ORAB and Sham-DB) were merged to a single Control group.

Differential gene expression analysis was performed using the DESeq2^62^ workflow (v1.46.0) and size factors calculated using the design formula ∼ *ConditionSimplified*. Genes with an FDR-adjusted p-value < 0.05 were called as differentially expressed. LFC shrinkage estimates were calculated using the apeglm method^63^ before gene ranking and visualisation. Log2FC for all contrasts were compared in a pairwise manner using Pearson’s correlation.

To identify groups of genes with similar trends in gene expression changes along the three profiled timepoints (Control, ORAB and DB), we first calculated the Euclidean distances between the average scaled and normalised counts per group for all DEGs (FDR < 0.05). Then, these distances were used to construct a hierarchical cluster that was cut to obtain 6 groups. This number was selected after visually inspecting the coherence of the gene expression trends and dendrogram branch heights for several resolutions (2-12 groups).

For the annotation of gene clusters, an overrepresentation analysis was run using decoupler^64^ v1.9.2 with ora method. Pathway and TF annotations were extracted from the CollecTRI^65^ and MSigDB mouse hallmarks^66^ databases through decoupler. Enrichment p-values were corrected using FDR and an adjusted p-value < 0.05 threshold was used for calling statistical significance.

### 4.13 snRNA-seq library preparation and sequencing

After cervical dislocation, mouse hearts were removed, left ventricles were isolated and flash frozen in liquid nitrogen. The LV samples were stored in -80°C. Isolation of nuclei was done with the 10X Genomics Chromium Nuclei Isolation kit including RNase Inhibitor (10X Genomics, USA, PN-1000494) according to the manufacturer’s protocol. Nuclei were stained with Ethidium Homodimer-2 (Invitrogen, USA) and counted with a fluorescent microscope (Keyence, Japan). SnRNA-seq libraries were generated using Chromium Single Cell 3′ Reagent Kits v3.1 (10x Genomics) following the manufacturer’s instructions. Approximately 10,000 nuclei were loaded per lane onto the Chromium microfluidic chips and processed following the kit protocol, including 11, 12, or 14 cycles of cDNA amplification and indexing PCR. Sequencing was performed on a NovaSeq 6000 (Illumina, USA) using the NovaSeq 6000 S4 Reagent Kit v1.5 (200 cycles) with paired-end reads (read 1: 28 bp, read 2: 90 bp; index 1: 10 bp, index 2: 10 bp). Raw sequencing data were processed using CellRanger (v7.0.0, 10× Genomics, USA) for demultiplexing, barcode counting, UMI counting, and feature-barcode matrix generation.

### 4.14 snRNA-seq quality control, integration and clustering

Individual samples were assessed to remove low quality nuclei using Scanpy^67^ v1.11.0 in Python v3.10.16. Nuclei with less than 300 genes and mitochondrial transcript proportion higher than 10% were excluded from further analysis. Genes expressed in less than 10 nuclei per sample were removed. The top 1% of nuclei with the largest number of genes with non-zero counts (n_genes_by_counts) were removed. Nuclei with a Scrublet^68^ v0.2.3 doublet score larger than 0.1 were called as doublets and excluded from further analysis.

For visualisation in a lower dimensionality space, samples were first aggregated and their counts were log-normalised. Highly variable genes were identified in a sample-specific fashion using Seurat’s method as implemented in Scanpy. Scaled counts for the 1,500 most variable genes shared by 75% of the samples (12 out of 16 samples) were used for Principal Component Analysis (PCA). Aggregated samples were visualised using the Uniform Manifold Approximation and Projection (UMAP). Then, samples were integrated by running Harmony^69^ using the technical batch as a covariate. An UMAP representation of the integrated data was visualised by computing the neighbourhood graph on 20 Harmony-corrected PCs. This nearest-neighbor graph was clustered using Leiden’s algorithm and resolution values of 0.4, 0.6, 0.8 and 1. A final resolution value of 0.6 was selected as it provided an appropriate level of cluster granularity upon visual inspection.

### 4.15 snRNA-seq annotation and identification of cell states

The snRNA-seq was annotated using 3 independent strategies, First, clusters were annotated by enriching the cell type markers defined in ReHeat2^21^ on the cluster marker genes. Cluster marker genes were calculated using Scanpy. Only marker genes with a logFC > 1.5 and a p-value < 0.001 were considered for this annotation. The enrichment was performed using hypergeometric tests. In parallel, the snRNA-seq data was annotated to all heart models^20,22,31,70,71^ available in CellTypist^27^. For annotation with CellTypist, raw counts were log-normalised and gene symbols in the models were translated from human to mouse. CellTypist was run with default mode “best match”, deactivated majority voting. Lastly, the expression of well characterised cell type markers was evaluated. Final cluster annotations into 9 major cell types (endothelial cells, fibroblasts, lymphatic endothelial cells (LECs), lymphoid cells, myeloid cells, neuronal cells, pericytes, ventricular cardiomyocytes (CM) and vascular smooth muscle cells (vSMCs)) were obtained from the consensus of the individual annotation strategies. Two clusters (clusters 20 and 21) were removed from further analysis, as they showed expression of marker genes from different cell types and neither the automatic annotation models nor the expression of known marker genes provided confident annotations. Cell cycle phase annotation was performed using scanpy’s score_genes_cell_cycle() function and Tirosh *et al*. G2M and S gene sets as reference^72^.

The identification of cellular states was conducted on CMs, endothelial cells and fibroblasts, as these were the cell types showing the largest number of DEGs across conditions. Each major cell type was subsetted and re-processed as before, setting a resolution of 0.4 for Leiden clustering. Clusters were annotated using literature-described marker genes and the Litvinukova *et al*^31^ CellTypist model.

### 4.16 snRNA-seq pseudobulks and compositional analysis

For the analysis of the cell type compositions and the identification of cell type-specific differentially expressed genes between conditions, pseudobulks were created using decoupler^64^ v1.9.2 by adding nuclei raw counts after grouping by sample and cell type. Pseudobulks with less than 10 nuclei or less than 1000 counts were filtered out. Neuronal pseudobulks were disregarded due to the low number of nuclei detected per sample. A PCA was performed on the scaled and log-normalised pseudobulk counts for quality control, and an ANOVA test was used to associate each PC with condition (Control, ORAB, DB), technical batch and cell type variables.

Compositional analysis was calculated for major cell types and cell states using ANOVA of the centered-log-ratio of cell proportions per sample. The p-values were Bonferroni-corrected for multiple testing. For cell types or states showing significant ANOVA results, a Tukey HSD test with Bonferroni-correction of p-values was applied to calculate pairwise differences in cell abundances.

### 4.17 snRNA-seq differential expression analysis and comparison with bulk DEGs

Differential gene expression was run using EdgeR^73^ on the pseudobulks. Three comparisons were calculated: ORAB vs Control (adverse remodeling), DB vs Control, and DB vs ORAB (reverse remodeling). P-values were adjusted using Benjamini-Hochberg. The mapping of cell type-specific DEGs on the bulk DEGs was conducted using decoupleR v.2.12.0 with the ulm method^64^.

### 4.18 Identification of multicellular programs

For the identification of multicellular programs, we fitted a Multicellular Factor Analysis model using MOFA2^74^ was used. For this, pseudobulk expression profiles calculated from at least 10 cells and a minimum of 50 genes were used. In each pseudobulk, genes with less than 20 counts in a single sample or detected in less than 40% of the remaining samples were discarded. The pseudobulk matrix for each cell type was normalized using the trimmed-mean of M values method in edgeR^73^ v4.0.2 with a scale factor of 1 million and log-transformed. Marker genes of other cell types were filtered out in the processed pseudobulk matrices to reduce the levels of contamination.

The Factor values were associated with clinical covariates using ANOVAs or linear models for categorical and continuous covariates, respectively. P-values were corrected across tests using the BH procedure.

Functional enrichment of the cell type-specific weights for each Factor was conducted using decoupleR v2.12.0 with ulm method^64^. Pathway and TF annotations were extracted from the CollecTRI^65^ and MSigDB mouse hallmarks^66^ databases through decoupler. Enrichment p-values for each database were corrected using the BH method.

For the projection the multicellular programs in the bulk clusters, we enriched the bulk cluster genes (weight = 1 if gene belongs to a cluster, else weight = 0) in the cell type gene-loadings of Factors 1 and 2 using decoupler^64^ v1.9.2 with the ulm method.

### 4.19 Ligand-receptor analysis

Ligand-receptor analysis was performed using LIANA+^32^ using differentially expressed genes calculated by edgeR and the snRNA-seq data as input. Differential gene expression results per gene were combined into statistics of potentially deregulated L-R interactions using LIANA+ multi.df_to_lr function using the mouse consensus resource. L-R interactions were filtered to remove lowly expressed interactions (interactions for which any of the genes are expressed in less than 10% of the nuclei).

To identify ligands that could be mediating CM recovery during reverse remodelling, the LIANA+ DB vs ORAB CCC results were filtered to (1) keep only interactions for which the ligand, receptor and interaction have FDR < 0.05; (2) keep only interactions for which the ligand are receptor logFC is overexpressed in DB vs ORAB (LogFC > 0); and (3) keep L-R interactions for which the ligand is a secreted protein. Human secreted vs non-secreted ligand annotations were obtained from Omnipath^75^ python client v.1.0.9 and translated to mouse gene symbols using HCOP database ortholog information. CCC networks were visualised using Cytoscape^76^ v3.10.3.

### 4.20 Identification of RELN target genes in cardiomyocytes

To predict target genes of RELN signaling in CMs, we used the NicheNet tool^36^ as implemented in the nichenetr R package v2.2.0 with the NicheNet v2 mouse prior model. NicheNet was run separately for each cell type that LIANA+ predicted to receive at least one significant interaction. For each receiver cell type, three inputs were defined: (1) Potential ligands: the candidate ligands that LIANA+ predicted to act on that cell type; (2) Background gene set: genes detected in at least 10% of the receiver’s cells and present in the ligand–target matrix; (3) Gene set of interest: DEGs in that cell type in the DB vs ORAB comparison (adj. p-value < 0.05). For each candidate ligand, predicted target genes were defined as the genes in the gene set of interest that were also among that ligand’s 200 top-ranked targets by regulatory potential (get_weighted_ligand_target_links, n = 200).

### 4.21 Statistical analysis

Statistical analyses of experimental and phenotypic data were performed using GraphPad Prism (version 10.6.1 [892]; GraphPad Software, USA). For comparisons among three or more independent groups, ordinary one-way analysis of variance (ANOVA) followed by Tukey’s multiple-comparisons test was used, as indicated in the respective figure legends. Paired circulating RELN measurements obtained before and after transcatheter aortic valve replacement were compared using the Wilcoxon matched-pairs signed-rank test. Data are presented as mean ± standard error of the mean (SEM), unless otherwise specified. Statistical tests were two-sided, and P < 0.05 was considered statistically significant.

Statistical analyses of transcriptomic data were performed using the computational methods described in the corresponding subsections. Briefly, differential gene expression in bulk RNA-seq and single-nucleus RNA-seq pseudobulk datasets was assessed using DESeq2 and edgeR, respectively, with false-discovery-rate correction for multiple testing. Differences in cell-type and cell-state proportions were evaluated using ANOVA followed by multiple-comparison-adjusted Tukey HSD tests, as detailed above. Adjusted P values < 0.05 were considered statistically significant for these analyses, unless otherwise indicated. The statistical test applied, exact sample size, definition of biological replicates and P values for individual comparisons are provided in the corresponding figure legends and supplementary information.

## 5. Code and data availability

The code used for data analysis is publicly accessible at https://github.com/saezlab/heart_revremod_paper.

All raw sequencing data used in this study will be uploaded and made publicly available upon publication. The bulk RNA-seq counts and metadata, as well as the processed snRNA-seq AnnData object are publicly available for download at https://doi.org/10.5281/zenodo.18214716.

## 6. Acknowledgements

This work was supported by Deutsche Forschungsgemeinschaft (DFG) (Collaborative Research Center CRC1550 “Molecular Circuits of Heart Disease”, INST 35/1699-1) to MD, BM, JB, JSR, and NF. MK and AYR are supported by DFG grants KU 4356/1-1 and RA 2717/5-1, respectively. We thank the Next Generation Sequencing Core Facility of the Medical Faculty Mannheim at Heidelberg University for single nuclei RNA-seq and preliminary data processing. The authors gratefully acknowledge data storage service SDS@hd and the support by the state of Baden-Württemberg through bwHPC and the German Research Foundation (DFG) through grants INST 35/1503-1 FUGG and INST 35/1597-1 FUGG respectively. We also thank S02 and INF projects of CRC1550 for disease modeling, biobanking and data management.

## 8. Authors contributions

MS, SN, MD, AYR, JSR, and NF conceived and designed the experiments; MS, IRG, RORF, AD, JP, SN, IA, MT, VA, KM, MD, MK and AYR performed the experiments and/or analyzed the data; VA, KM, BM, FL, MK, JB, AYR, JSR and NF contributed reagents/materials/analysis tools; MS, IRG and AYR wrote the manuscript; RORF, BM, MK, JSR and NF revised the manuscript. All authors have read and agreed to the submitted manuscript.

## Supplementary Materials

### Supplementary Files

***Supplementary File 1****. Bulk RNA-seq DEGs for the ORAB vs DB, ORAB vs Control and DB vs Control contrasts*.

***Supplementary File 2****. Hierarchical cluster membership of bulk RNA-seq DEGs*.

***Supplementary File 3****. Functional enrichment results of each bulk RNA-seq cluster*.

***Supplementary File 4****. snRNA-seq cluster marker genes*.

***Supplementary File 5****. Differentially expressed genes for each cell type across conditions, calculated with snRNA-seq pseudobulks*.

***Supplementary File 6****. Multicellular factors functional enrichment*.

***Supplementary File 7****. Marker genes of the CM cell states*.

***Supplementary File 8****. Marker genes of the endothelial cell states*

***Supplementary File 9****. Marker genes of the fibroblast cell states*

***Supplementary File 10****. Deregulated CCC interactions in ORAB vs. Control*

***Supplementary File 11****. Deregulated CCC interactions in DB vs. ORAB*.

***Supplementary File 12****. Candidate CCC interactions driving reverse remodelling*.

***Supplementary File 13****. List of primers used in RT-qPCR experiments*.

### Supplementary Figures

**Figure S1.**
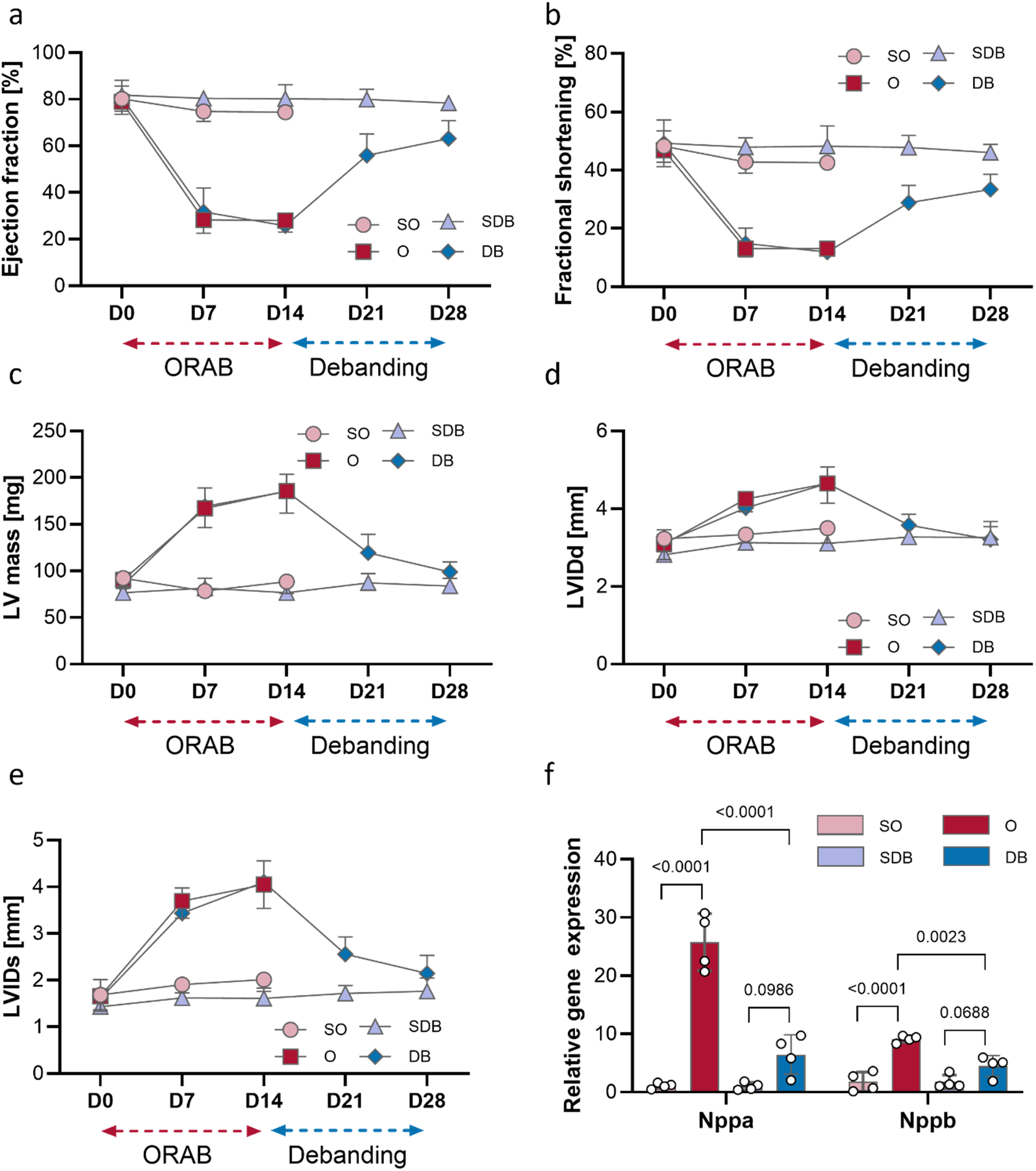
Reversal of pressure overload induces partial functional and structural cardiac recovery. **a-b)** Quantification of left ventricular fractional shortening (LVFS) and left ventricular ejection fraction (LVEF) by echocardiography. ORAB resulted in marked systolic dysfunction, whereas DB significantly improved cardiac contractile performance, although values did not fully normalize to sham levels. LVFS and LVEF were indistinguishable between SO and SDB. **c)** Left ventricular mass assessed by echocardiography. ORAB markedly increased LV mass, which was significantly reduced after unloading by de-banding. No differences were observed between time-matched sham cohorts. **d-e)** Left ventricular internal diameter at diastole (LVIDd) and systole (LVIDs). Pressure overload induced pathological chamber dilation that was partially reversed following de-banding. Measurements were equivalent between sham groups euthanized at two or four weeks. For a-e, n = 10 for SO, 15 for O, 7 for SDB and 7 for DB. **f)** Quantitative PCR analysis of cardiac fetal gene expression. Transcripts encoding atrial natriuretic peptide (*Nppa*) and brain natriuretic peptide (*Nppb*) were robustly upregulated in ORAB hearts and significantly reduced following de-banding, although expression levels remained modestly elevated relative to sham controls. Gene expression was comparable between the two sham cohorts. n = 4 each per group. Data are presented as mean ± SEM. Statistical significance was assessed using ordinary one-way ANOVA followed by Tukey’s multiple comparisons test. Exact p-values for the respective comparisons are indicated in the graph.

**Figure S2.**
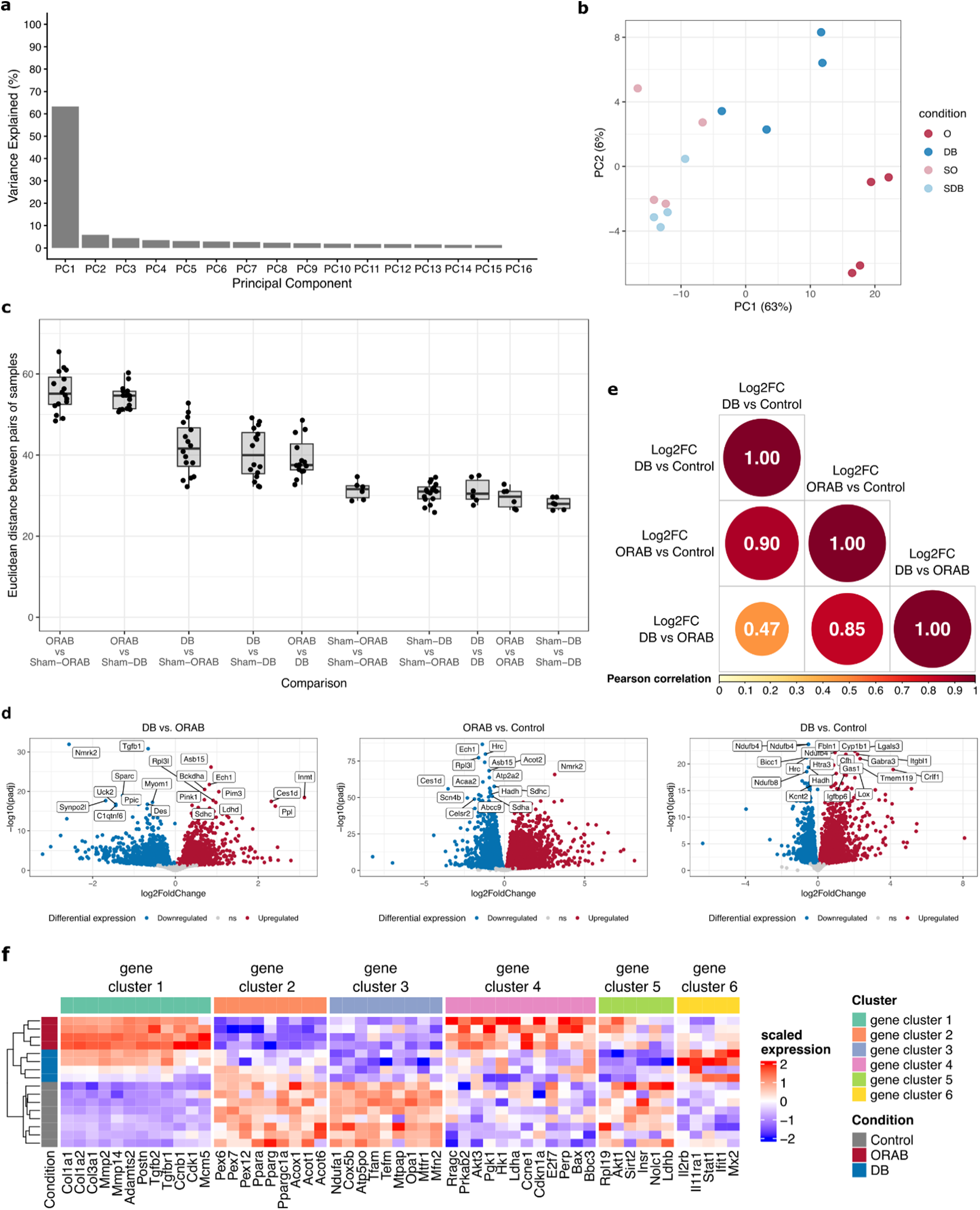
Bulk RNA-seq. **a)** Percentage of explained variance per PC for the bulk RNA-seq PCA. **b)** PCA visualisation of the bulk RNA-seq. Each point in the scatter-plot represents a biological sample. **c)** Boxplot of the euclidean distances between pairs of samples calculated based on their bulk mRNA expression levels. Each point represents the euclidean distance between a pair of samples. d) Volcano plots of the ORAB vs DB, ORAB vs Control and DB vs Control for the bulk RNA-seq. Colored genes have a FDR-adjusted p-value < 0.05 and a log2FC < 0 (blue) or log2FC > 0 (red). The top 20 most significant DEGs appear labeled. **e)** Heatmap of the Pearson correlation between Log2FC for DEGs between the ORAB vs DB, ORAB vs Control and DB vs Control contrasts. **f)** Heatmap of the scaled expression of selected genes representative of the biological function of each cluster.

**Figure S3.**
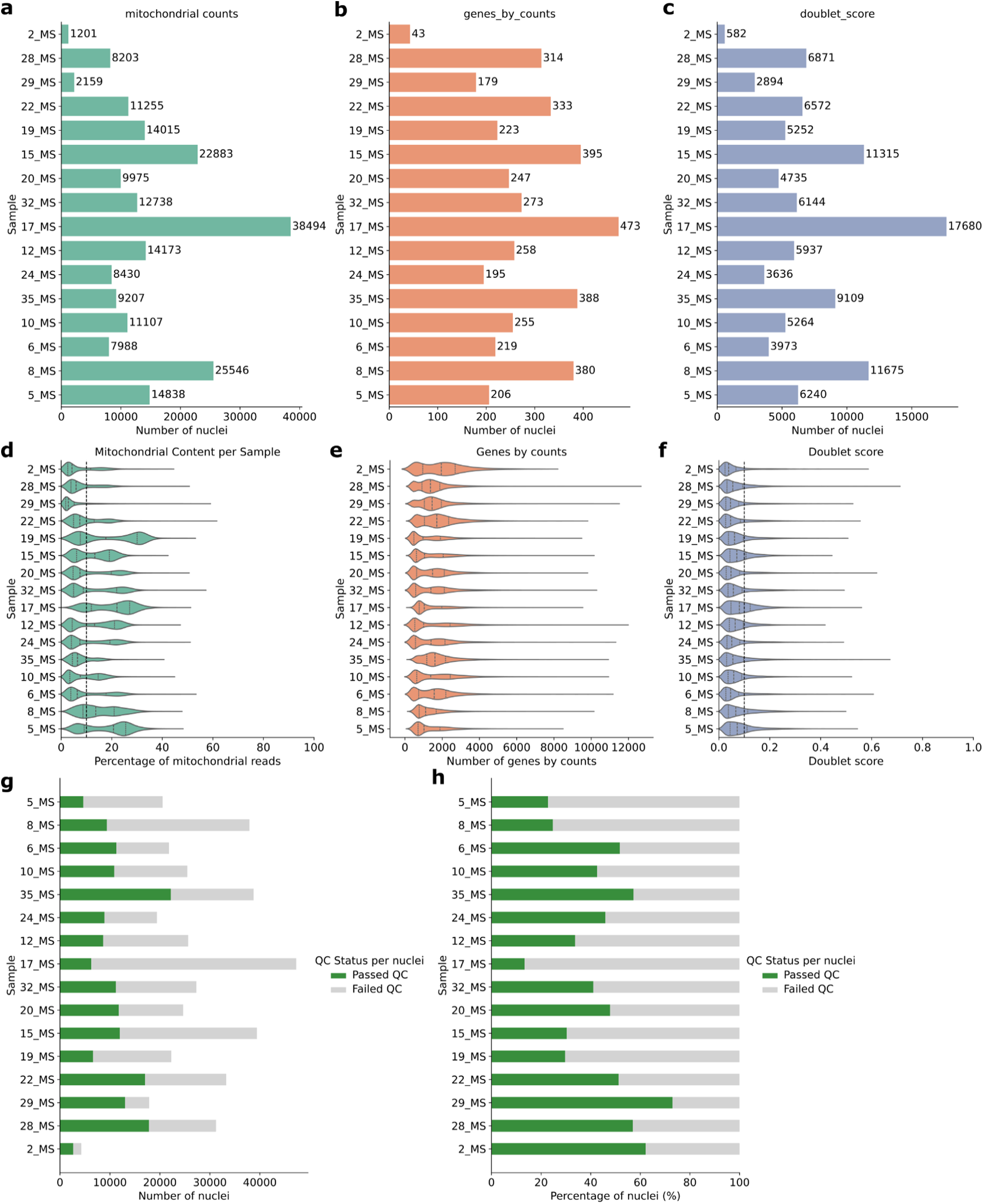
snRNA-seq quality control. **a-c)** Barplot of the number of nuclei per sample that did not pass a) mitochondrial count QC filter, **b)** genes_by_counts QC filter and **c)** Scrublet doublet score filter. **d)** Violin plot of the distribution of the percentage of mitochondrial reads per sample. The vertical line indicates the 10% threshold used for QC filtering. **e)** Violin plot of the distribution of gene_by_count values per sample. **f)** Distribution of the doublet score per sample. The vertical line indicates the 0.1 threshold used for QC filtering. **g)** Number of nuclei per sample that passed (green) or did not pass (light gray) the QC. **h)** Percentage of nuclei per sample that passed (green) or did not pass (light gray) the QC.

**Figure S4.**
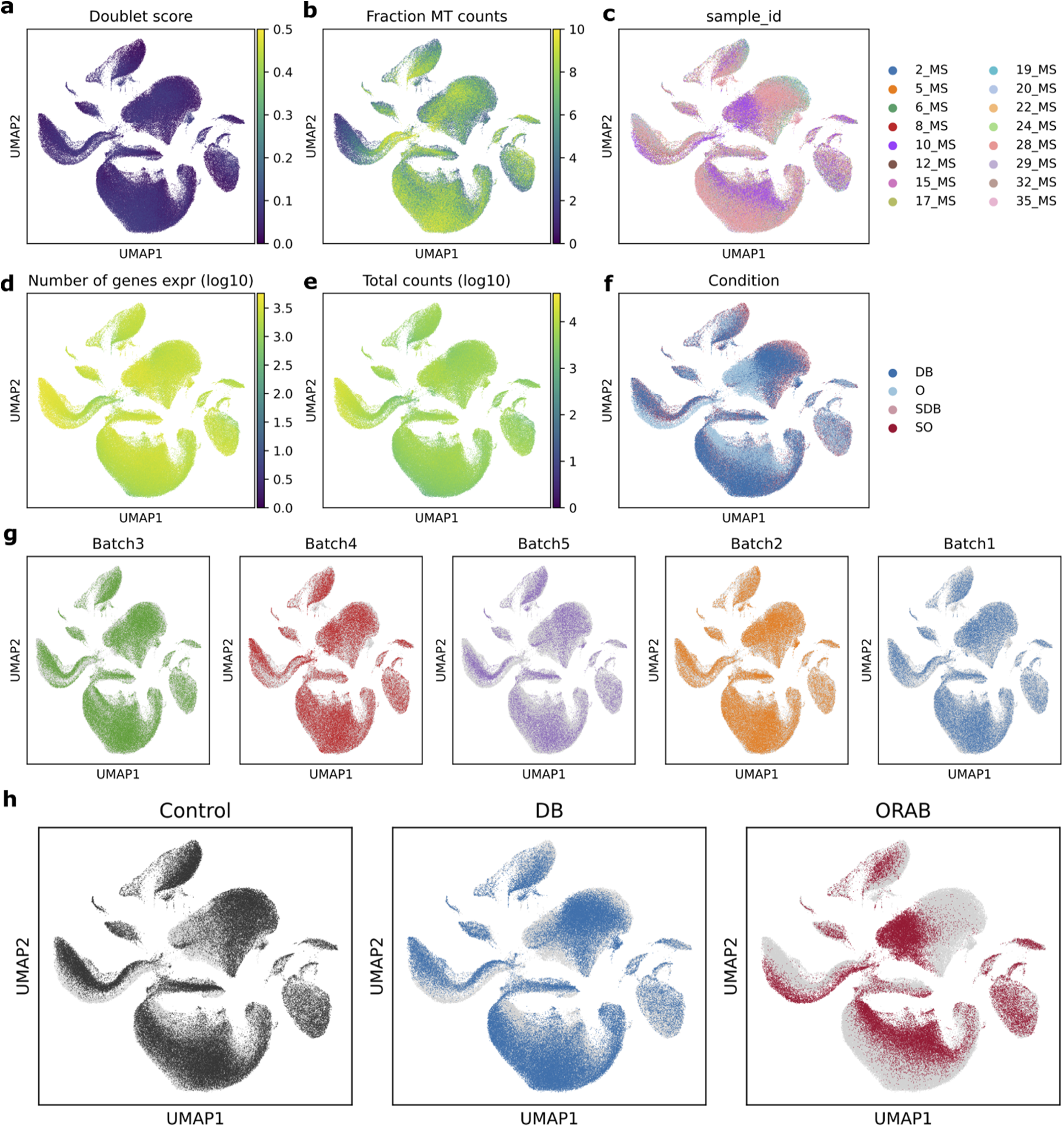
snRNA-seq data integration QC. **a-h)** UMAP visualisation of all integrated nuclei colored by **a)** the doublet score, **b)** fraction of mitochondrial counts, **c)** sample ID, **d)** number of expressed genes by counts, **e)** total counts, **f)** condition (O: ORAB, DB: DB, SO: Sham-ORAB, SDB: Sham-DB), **g)** technical batch and **h)** simplified condition.

**Figure S5.**
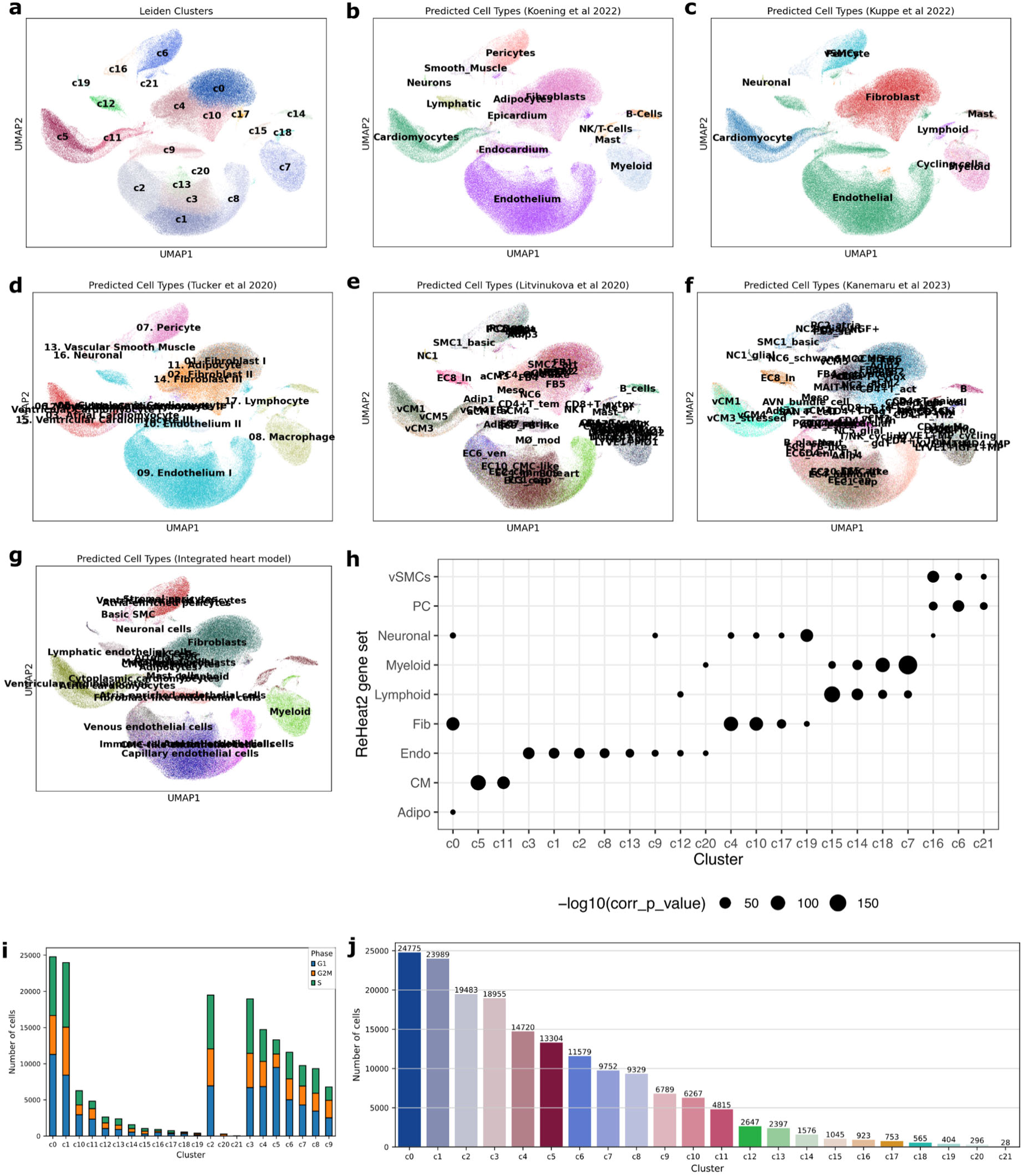
snRNA-seq clustering. **a)** UMAP visualisation of integrated nuclei colored by cluster. **b-g)** UMAP visualisation of integrated nuclei colored by the consensus annotation predicted by CellTypist using the **b)** Koenig *et al.* 2022 model, **c)** Kuppe *et al.* 2022 model, **d)** Tucker *et al.* 2020 model, **e)** Litvinukova *et al.*^31^ model, **f)** Kanemaru *et al.* 2023 model and **g)** the integrated heart model. **h)** Dotplot of the p-values obtained by enrichment analysis of ReHeat2^21^ cell type markers on cluster marker genes, calculated using hypergeometric tests. **i)** Barplot of the number of cells in each cell cycle phase (G1, S or G2M) per cluster). **j)** Barplot of the number of cells per cluster.

**Figure S6.**
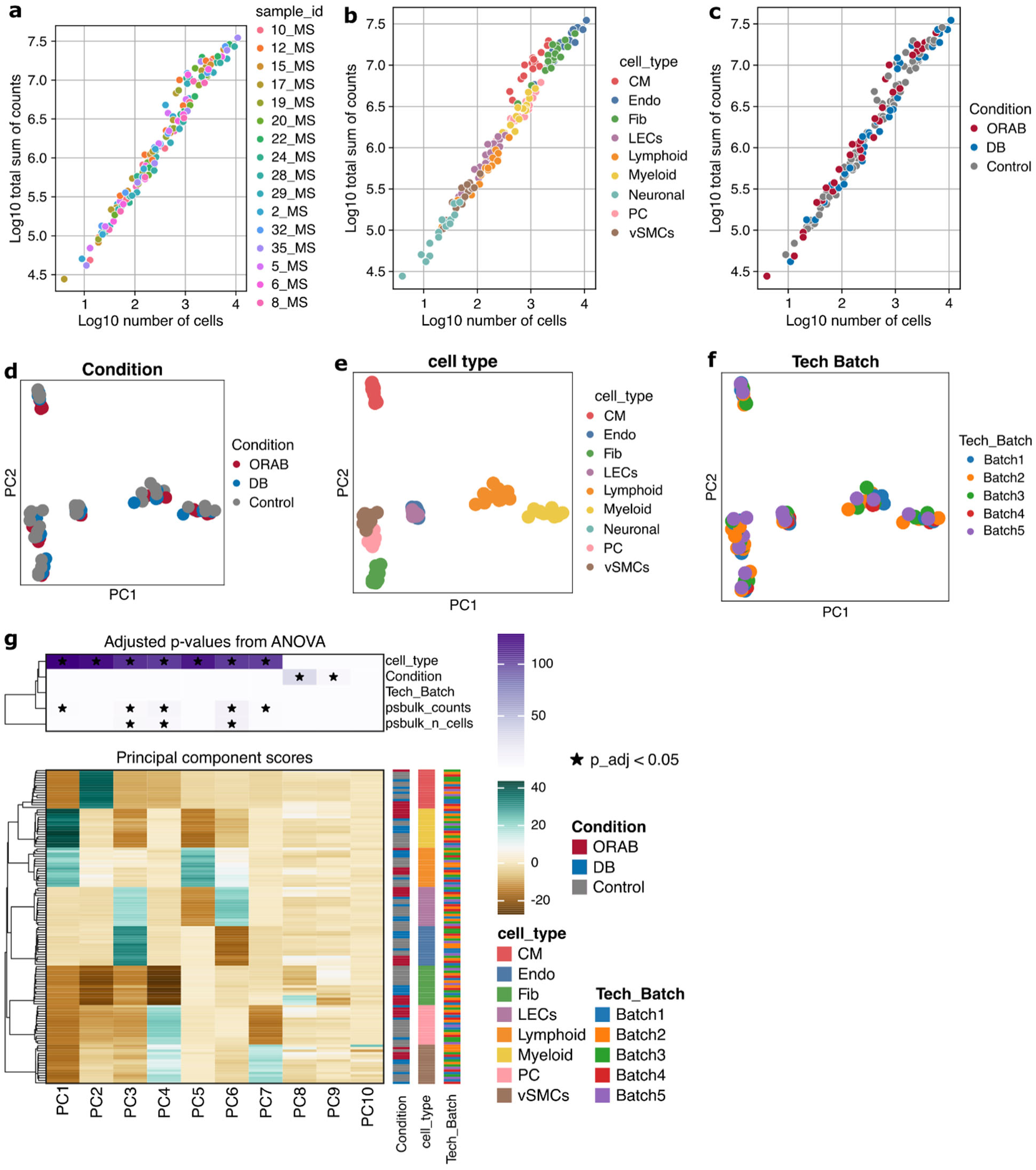
Pseudobulk QC. **a-c)** Scatterplot of the log10(total counts) and Log10(number of cells) in each pseudobulk, colored by **a)** sample ID, **b)** cell type and **c)** condition. **d-f)** PCA visualisation of the pseudobulks, colored by **d)** condition, **e)** cell type and **f)** technical batch. **g)** Heatmap of the PC loadings per pseudobulk and the ANOVA association between PCs and metadata variables. Stars indicate ANOVA Benjamini-Hochberg adjusted p-value < 0.05.

**Figure S7.**
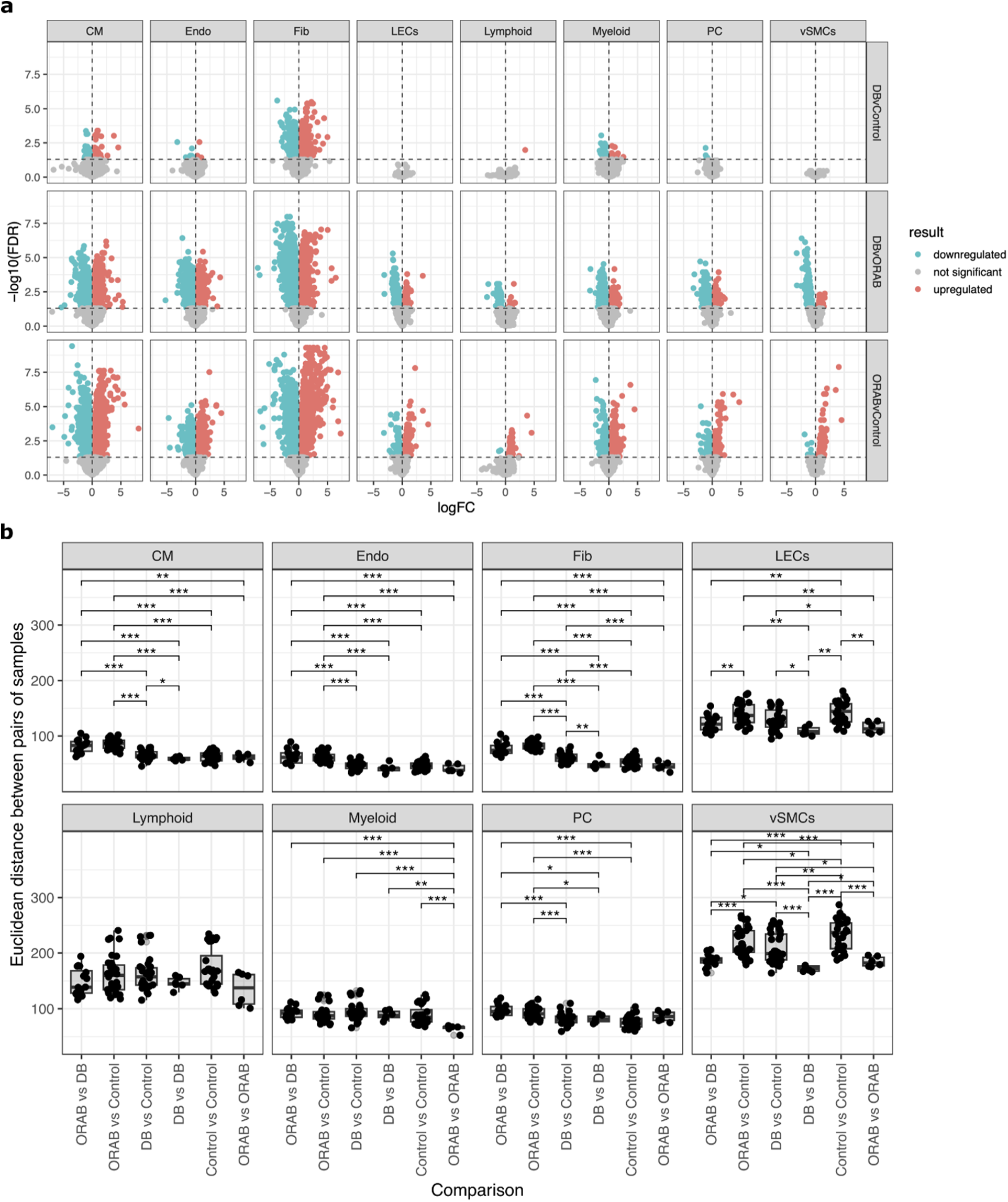
Cell type-specific differential expression analysis. **a)** Volcano plots of the differential expression analysis for each cell type. **b)** Boxplot of the euclidean distances between pairs of samples calculated based on their pseudobulk expression levels. Each point represents the euclidean distance between a pair of samples.

**Figure S8.**
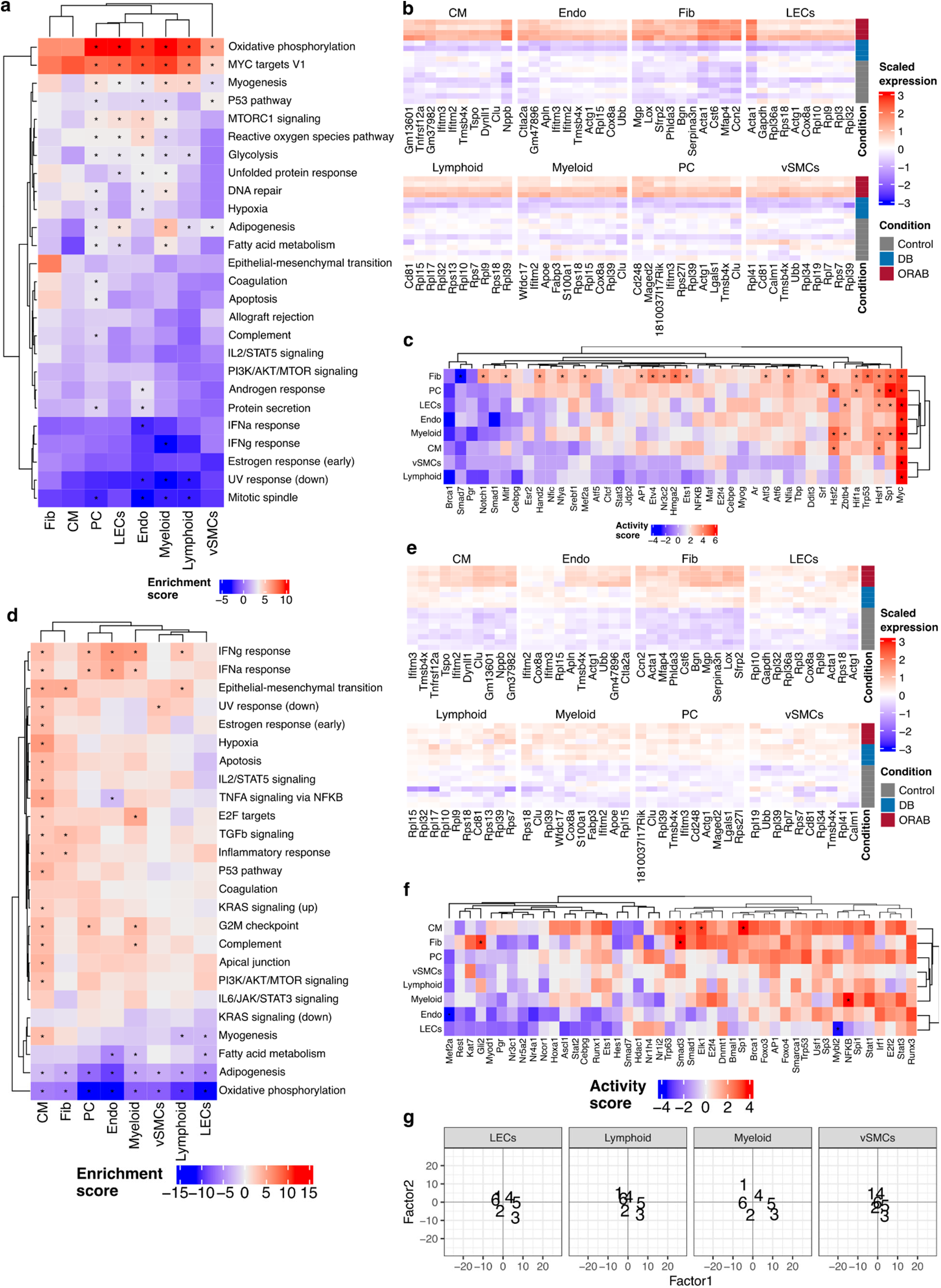
Functional enrichment of multicellular factors. **a)** Heatmap of the enrichment scores of the MSigDB Hallmarks pathways on the multicellular Factor1 cell type-gene loadings. Asterisks indicate FDR < 0.05. **b)** Heatmap of the scaled log-normalised expression of the top 10 genes with highest weights for Factor 1 per cell type. **c)** Heatmap of the activity scores of Collectri TFs on the multicellular Factor1 cell type-gene loadings. Asterisks indicate FDR < 0.05. Only TFs with |activity| > 2 are shown. **d)** Heatmap of the enrichment scores of the MSigDB Hallmarks pathways on the multicellular Factor2 weights. **e)** Heatmap of the scaled log-normalised expression of the top 10 genes with highest weights for Factor 2 per cell type. **f)** Heatmap of the activity scores of Collectri TFs on the multicellular Factor1 cell type-gene loadings. Asterisks indicate FDR < 0.05. Only TFs with |activity| > 2 are shown. **g)** Scatterplot of the enrichment scores of the gene clusters identified in bulk in the cell type-gene loadings of multicellular factors 1 and 2. Only the cell types not shown in Fig. 2h are shown.

**Figure S9.**
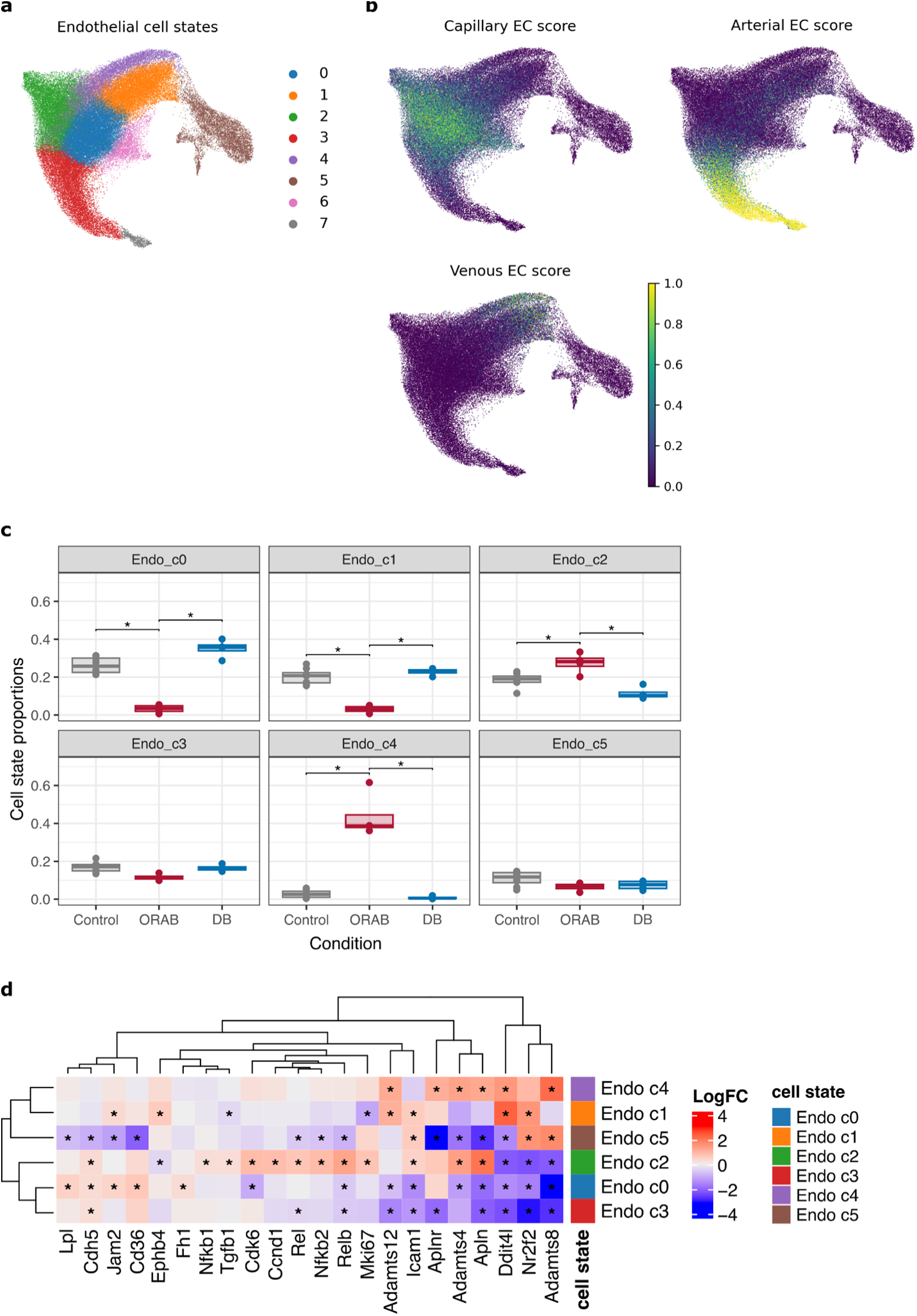
Endothelial cell states. **a)** UMAP visualisation of the endothelial cell states. **b)** UMAP visualisation of the endothelial cells, colored by the CellTypist score of similarity with the capillary, arterial and venous endothelial cell populations described by Litvinukova *et al.*^31^. **c)** Boxplots of the cell type proportions for the endothelial cell states. Each dot represents one biological replicate. Asterisks represent Tukey HSD adj. p-value < 0.05, for contrasts with ANOVA adj. P-value < 0.05. **d)** Heatmap of the LogFC of selected marker genes in the main endothelial cell states. The logFC were calculated using all other CMs as reference. Asterisks indicate FDR < 0.05.

**Figure S10.**
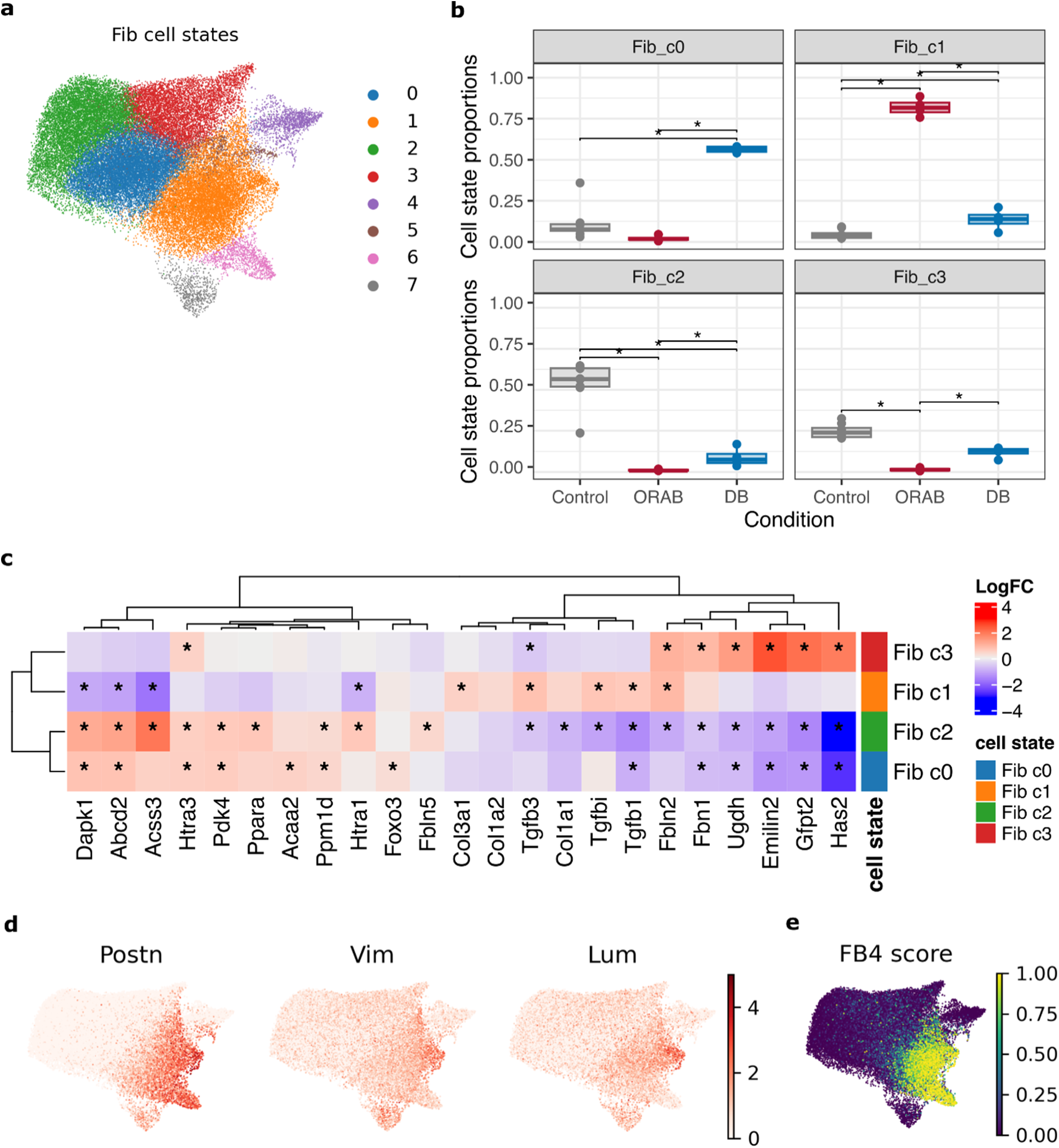
Fibroblast cell states. **a)** UMAP visualisation of the fibroblast cell states. **b)** Boxplots of the cell type proportions for the fibroblast cell states representing > 5% of all fibroblasts. Each dot represents one biological replicate. Asterisks represent Tukey HSD adj. p-value < 0.05, for contrasts with ANOVA adj. P-value < 0.05. **c)** Heatmap of the LogFC of selected marker genes in the main fibroblasts cell states. The logFC were calculated using all other fibroblasts as reference. Asterisks indicate FDR < 0.05. **d)** UMAP visualisation of the fibroblasts, colored by their log-normalised expression of the activated fibroblast marker genes *Postn*, *Vim* and *Lum*. **e)** UMAP visualisation of the fibroblasts, colored by the CellTypist score of similarity with the FB4 population of TGFb-responsive fibroblasts described by Litvinukova et al32.

**Figure S11.**
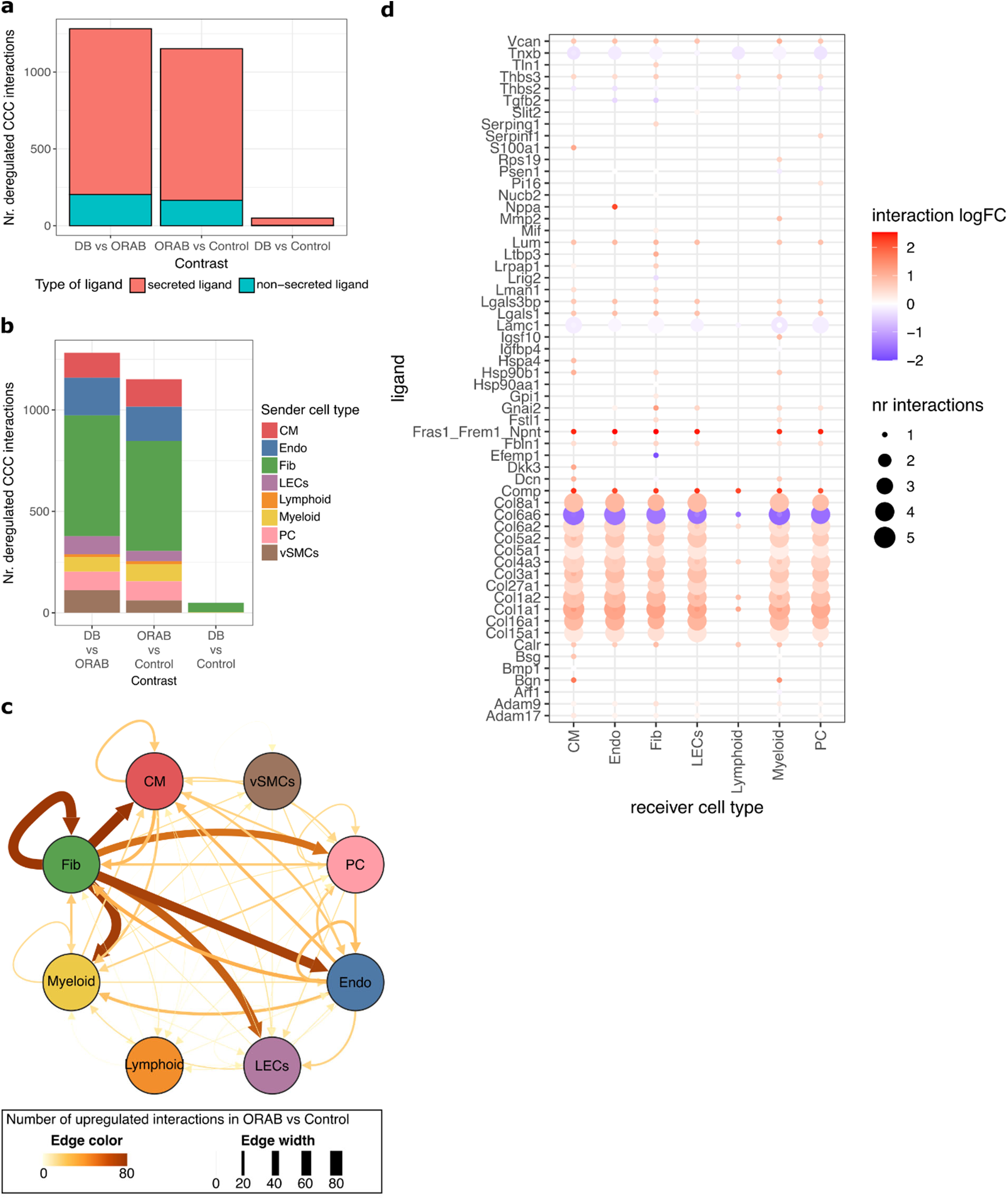
Cell-cell communication networks in adverse cardiac remodelling. **a)** Barplot of the number of CCC interactions detected by LIANA+, colored by type of ligand (secreted or non-secreted). **b)** Barplot of the number of CCC interactions detected by LIANA+, colored by the sender cell type. **c)** Network visualisation of the CCC interactions upregulated in ORAB vs Control. Edge color and width represent the number of interactions. **d)** Dotplot of the fibroblast ligands deregulated in ORAB vs Control. The dot color represents ligand logFC. The dot size represents the number of interactions mediated by the ligand.

**Figure S12.**
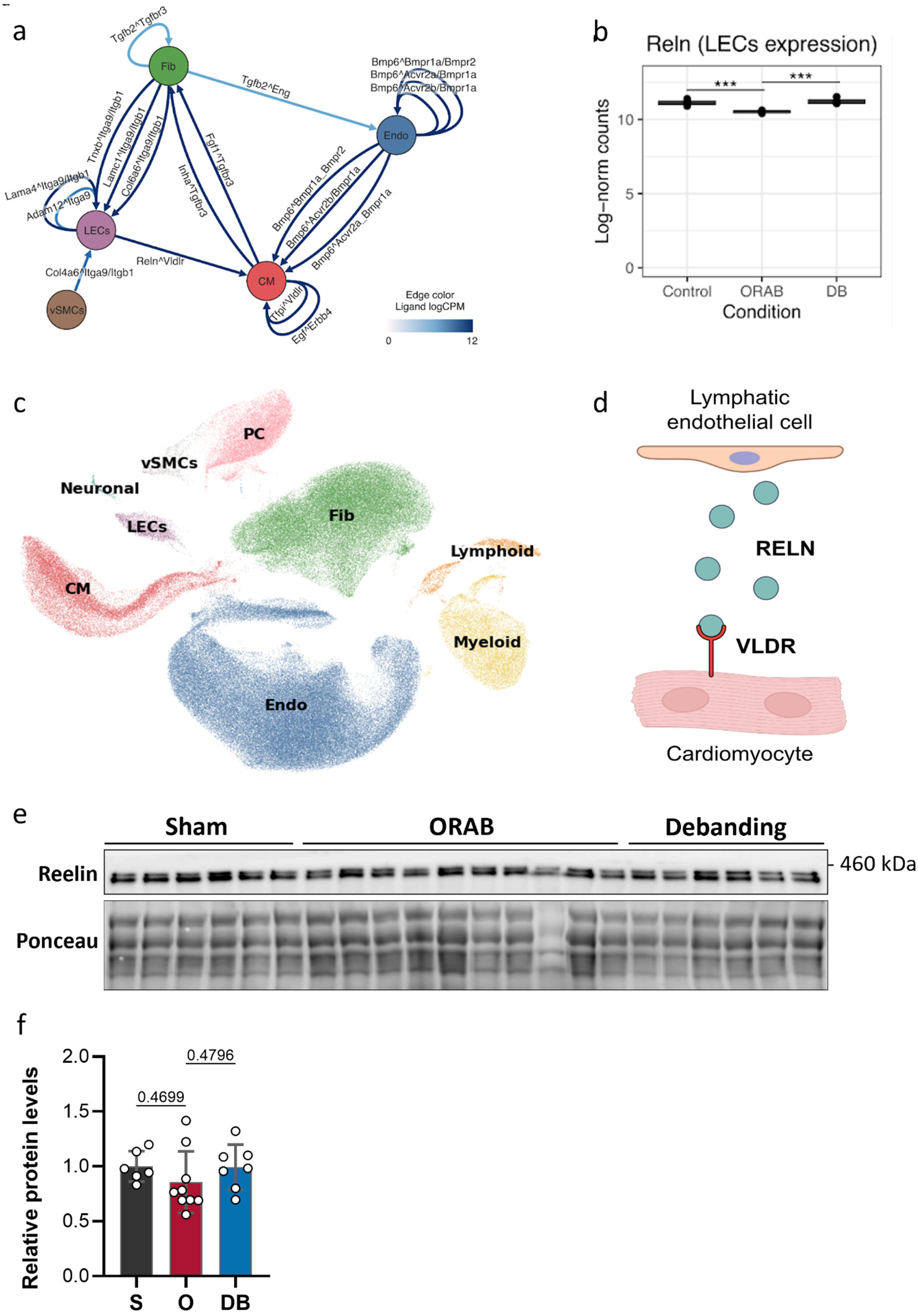
Identification, cellular localization, and translational relevance of RELN signaling during reverse cardiac remodeling. **a)** Cell-cell communication network highlights candidate intercellular signaling interactions associated with reverse remodeling. The analysis identified Reln-Vldlr as a candidate interaction between lymphatic endothelial cells (LECs) and cardiomyocytes (CMs). Nodes denote cardiac cell populations and edges indicate predicted ligand-receptor interactions; edge color reflects ligand expression as indicated. **b)** Pseudobulk expression of Reln in LECs from Control, ORAB, and DB hearts, showing suppression during pressure overload and restoration following de-banding. Each point represents a biological replicate. Asterisks indicate FDR < 0.001 (***). **c)** UMAP representation of the single-nucleus RNA-sequencing dataset showing the major cardiac cell populations used to contextualize the expression distributions shown in Fig. 5b. **d)** Schematic model of the proposed paracrine signaling interaction in which RELN is secreted by cardiac LECs binds VLDLR expressed on cardiomyocytes. **e)** Representative assessment of circulating RELN in plasma from Control, ORAB, and DB mice [insert exact assay/blot description here]. **f)** Quantification of plasma RELN abundance in Control, ORAB, and DB mice. n = 6 mice for sham, 9 for ORAB and 6 for de-banding. Data are presented as individual biological replicates as mean ± SEM. Statistical significance was assessed using ordinary one-way ANOVA followed by Tukey’s multiple comparisons test. Exact p-values for the respective comparisons are indicated in the graph. Control, sham (also denoted as ‘S’)-operated control; ORAB (also denoted as ‘O’), O-ring aortic banding; DB, de-banding. Fib, fibroblasts; Endo, endothelial cells; LECs, lymphatic endothelial cells; CM, cardiomyocytes; vSMCs, vascular smooth muscle cells; PC, pericytes; ORAB, O-ring aortic banding; DB, de-banding.

**Supplementary Figure S13.**
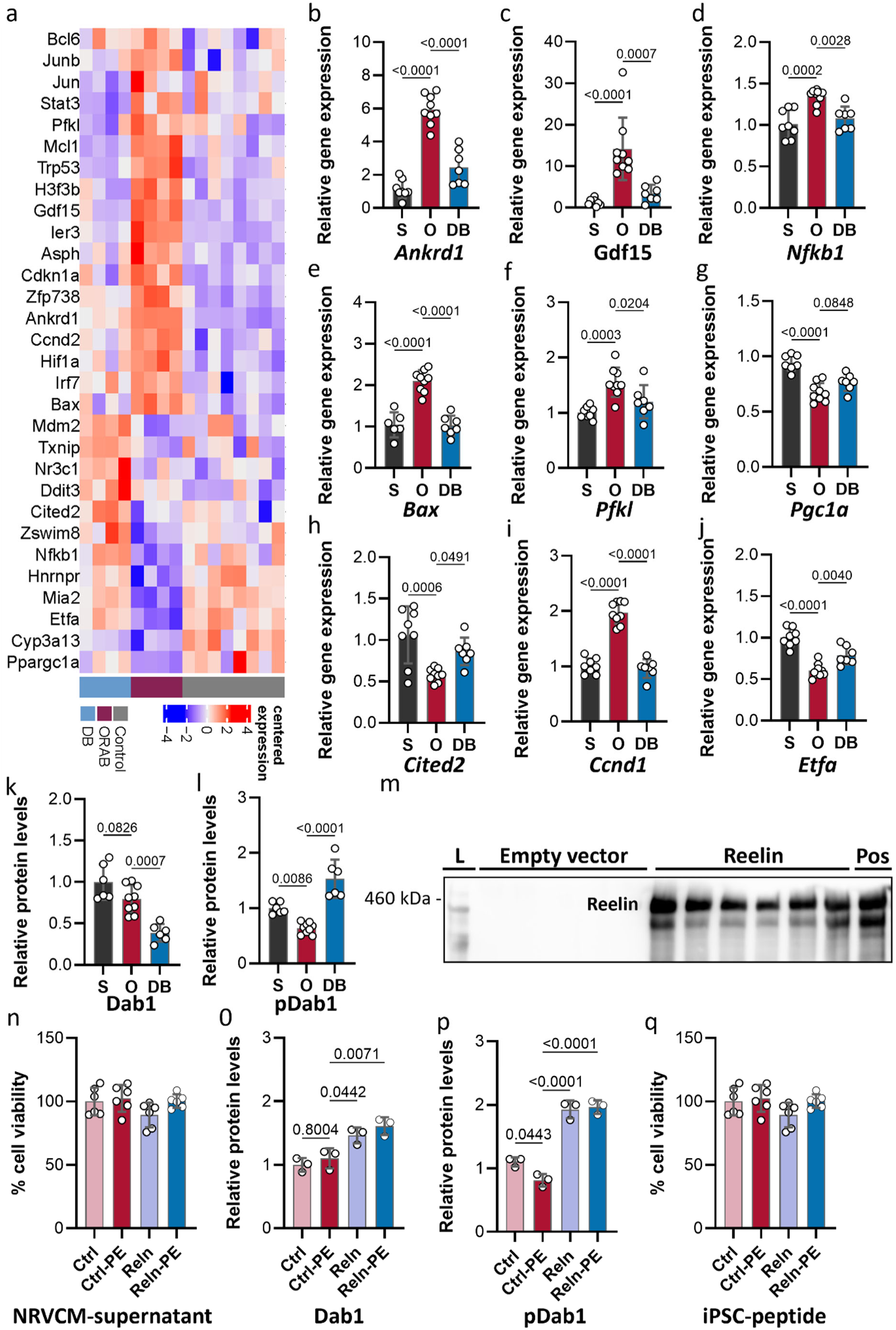
Transcriptional and biochemical characterization of RELN–VLDLR–DAB1 signaling during reverse cardiac remodeling and cardiomyocyte hypertrophic stress. **a)** Heatmap showing the expression of computationally predicted genes associated with RELN–VLDLR signaling across Control, ORAB and DB hearts. Expression values are centered and scaled across samples as indicated by the color scale. Experimental groups are indicated below the heatmap. **b–j)** Independent qPCR validation of nine selected candidate genes associated with the predicted RELN–VLDLR transcriptional program in cardiac tissue from Control, ORAB and DB mice, including *Ankrd1, Gdf15, Nfkb1, Bax, Pfkl, Ppargc1a, Cited2, Ccnd1* and *Etfa*. n = 8 mice for sham, 9 for ORAB and 7 for de-banding. **k,l)** Densitometric quantification of total DAB1 (**k**) and phosphorylated DAB1 (pDAB1; **l**) abundance in myocardial tissue from Control, ORAB and DB mice. Representative immunoblots are shown in Fig. 6a. n = 6 mice for sham, 9 for ORAB and 6 for de-banding. **m)** Representative immunoblot of culture supernatants collected from HEK293T cells transfected with empty vector (EV) or a RELN expression construct, confirming enrichment of secreted RELN in conditioned medium generated from RELN-overexpressing cells. **n)** MTT-based assessment of NRVCM viability following exposure to EV- or RELN-enriched conditioned medium under basal conditions or following PE-induced hypertrophic stress. n = 6 each biological replicates per group **o,p)** Densitometric quantification of total DAB1 (**o**) and pDAB1 (**p**) abundance in NRVCMs exposed to EV- or RELN-enriched conditioned medium under basal and PE-stimulated conditions. Representative immunoblots are shown in Fig. 6f. n = 3 biological replicates per group. **q)** MTT-based assessment of cell viability in human induced pluripotent stem cell-derived cardiomyocytes (hiPSC-CMs) treated with the biologically active RELN fragment under basal conditions or following phenylephrine (PE)-induced hypertrophic stress. n = 6 biological replicates per group. Data are presented as individual biological replicates with mean ± SEM. Statistical significance was determined as indicated in the respective panels and Methods. Control, sham-operated control; ORAB, O-ring aortic banding; DB, de-banding; EV, empty vector; RELN, Reelin; VLDLR, very-low-density lipoprotein receptor; DAB1, Disabled-1; PE, phenylephrine; NRVCM, neonatal rat ventricular cardiomyocyte; Pos = positive control, mouse brain protein lysate.

### Supplementary Tables

**Table S1.**
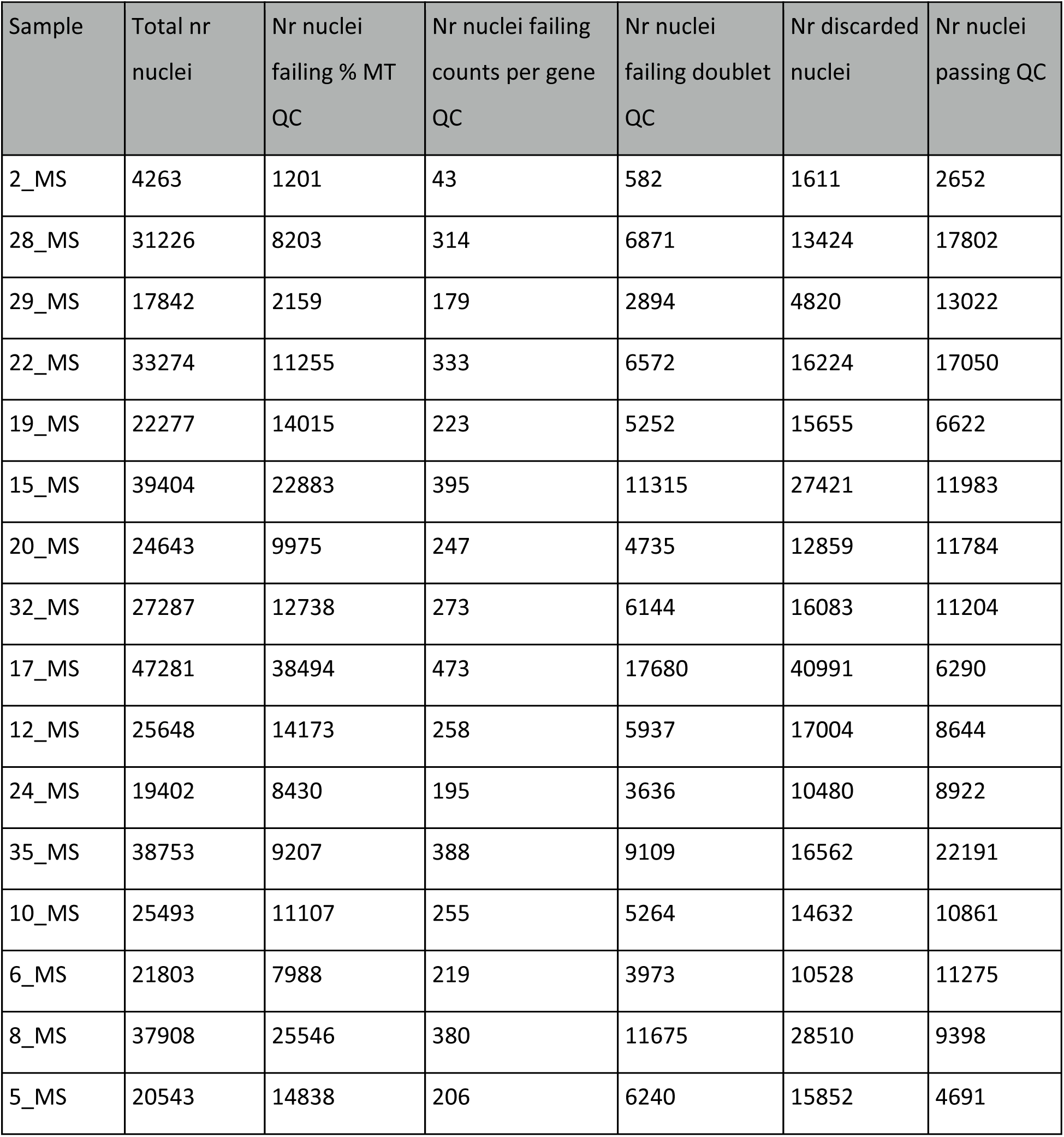
Number of nuclei passing and failing QC per sample.

**Table S2.** ANOVA results for the compositional analysis of major cell types.

| cell_type | term | df | sumsq | meansq | statistic | p-value | adj. p-value |
|---|---|---|---|---|---|---|---|
| CM | Condition | 2 | 0.192 | 0.096 | 3.896 | 4.725e-02 | 7.560e-02 |
| Endo | Condition | 2 | 0.351 | 0.176 | 16.923 | 2.406e-04 | 6.416e-04 |
| Fib | Condition | 2 | 0.212 | 0.106 | 6.726 | 9.877e-03 | 1.975e-02 |
| LECs | Condition | 2 | 0.139 | 0.070 | 1.538 | 2.515e-01 | 3.289e-01 |
| Lymphoid | Condition | 2 | 0.351 | 0.175 | 1.251 | 3.185e-01 | 3.289e-01 |
| Myeloid | Condition | 2 | 1.090 | 0.545 | 25.943 | 2.895e-05 | 1.158e-04 |
| PC | Condition | 2 | 1.932 | 0.966 | 91.134 | 2.247e-08 | 1.797e-07 |
| vSMCs | Condition | 2 | 0.175 | 0.088 | 1.2127 | 3.289-01 | 3.289e-01 |

**Table S3.** Multicellular program model R2 values for the fit of each Factor to each cell type.

| Cell type | Factor1 R2 | Factor2 R2 | Factor3 R2 | Factor4 R2 |
|---|---|---|---|---|
| CM | 0.490 | 0.119 | 0.052 | 0.016 |
| Endo | 0.547 | 0.058 | 0.094 | 0.015 |
| Fib | 0.514 | 0.149 | 0.084 | 0.008 |
| LECs | 0.424 | 0.075 | 0.080 | 0.065 |
| Lymphoid | 0.219 | 0.048 | 0.060 | 0.160 |
| Myeloid | 0.355 | 0.094 | 0.082 | 0.064 |
| PC | 0.484 | 0.066 | 0.105 | 0.024 |
| vSMCs | 0.330 | 0.066 | 0.064 | 0.105 |
| Mean | 0.420 | 0.084 | 0.078 | 0.057 |

**Table S4.** ANOVA results for the compositional analysis of CM cell states.

| Cluster | Nr nuclei | % of all CM nuclei | term | df | sumsq | meansq | statistic | p-value | adj. p-value |
|---|---|---|---|---|---|---|---|---|---|
| CM_c0 | 8,181 | 45.15 | Condition | 2 | 24.781 | 12.390 | 48.605 | 9.251e-07 | 9.251e-06 |
| CM_c1 | 4,235 | 23.37 | Condition | 2 | 1.084 | 0.542 | 1.734 | 0.215 | 0.430 |
| CM_c2 | 3,785 | 20.89 | Condition | 2 | 11.022 | 5.511 | 11.313 | 0.001 | 0.007 |
| CM_c3 | 670 | 3.7 | Condition | 2 | 0.457 | 0.228 | 0.773 | 0.482 | 0.688 |
| CM_c4 | 448 | 2.47 | Condition | 2 | 1.040 | 0.520 | 2.298 | 0.140 | 0.430 |
| CM_c5 | 308 | 1.7 | Condition | 2 | 1.308 | 0.654 | 1.906 | 0.188 | 0.430 |
| CM_c6 | 201 | 1.11 | Condition | 2 | 0.120 | 0.060 | 0.109 | 0.898 | 0.898 |
| CM_c7 | 126 | 0.70 | Condition | 2 | 1.232 | 0.616 | 1.483 | 0.263 | 0.438 |
| CM_c8 | 100 | 0.55 | Condition | 2 | 0.386 | 0.193 | 0.305 | 0.743 | 0.825 |
| CM_c9 | 65 | 0.36 | Condition | 2 | 0.288 | 0.144 | 0.370 | 0.698 | 0.825 |

**Table S5.** ANOVA results for the compositional analysis of endothelial cell states.

| Cluster | Nr nuclei | % of all Endo nuclei | term | df | sumsq | meansq | statistic | p-value | adj. p-value |
|---|---|---|---|---|---|---|---|---|---|
| Endo_c0 | 20,947 | 25.88 | Condition | 2 | 14.913 | 7.457 | 42.988 | 1.861e-06 | 7.443e-06 |
| Endo_c1 | 14,551 | 17.98 | Condition | 2 | 11.070 | 5.535 | 43.773 | 1.680e-06 | 7.443e-06 |
| Endo_c2 | 14,214 | 17.56 | Condition | 2 | 2.059 | 1.029 | 17.296 | 2.171e-04 | 4.342e-04 |
| Endo_c3 | 12,559 | 15.52 | Condition | 2 | 0.149 | 0.074 | 1.784 | 0.208 | 0.276 |
| Endo_c4 | 7,750 | 9.57 | Condition | 2 | 43.100 | 21.550 | 30.078 | 1.328e-05 | 3.540e-05 |
| Endo_c5 | 6,950 | 8.59 | Condition | 2 | 0.147 | 0.074 | 0.534 | 0.598 | 0.683 |
| Endo_c6 | 3,060 | 3.78 | Condition | 2 | 0.582 | 0.291 | 2.675 | 0.106 | 0.170 |
| Endo_c7 | 911 | 1.13 | Condition | 2 | 0.200 | 0.100 | 0.271 | 0.767 | 0.767 |

**Table S6.** ANOVA results for the compositional analysis of fibroblast cell states.

| Cluster | Nr nuclei | % of all Fib nuclei | term | df | sumsq | meansq | statistic | p-value | adj. p-value |
|---|---|---|---|---|---|---|---|---|---|
| Fib_c0 | 12,389 | 26.63 | Condition | 2 | 20.509 | 10.254 | 22.233 | 6.375e-05 | 1.275e-04 |
| Fib_c1 | 12,070 | 25.95 | Condition | 2 | 38.415 | 19.207 | 87.920 | 2.793e-08 | 2.234e-07 |
| Fib_c2 | 10,802 | 23.22 | Condition | 2 | 24.665 | 12.332 | 37.884 | 3.775e-06 | 1.007e-05 |
| Fib_c3 | 6,770 | 14.55 | Condition | 2 | 13.4711 | 6.736 | 40.564 | 2.579e-06 | 1.007e-05 |
| Fib_c4 | 1,473 | 3.17 | Condition | 2 | 2.060 | 1.030 | 1.645 | 0.231 | 0.231 |
| Fib_c5 | 1,213 | 2.61 | Condition | 2 | 2.805 | 1.403 | 15.345 | 3.786e-04 | 5.048e-04 |
| Fib_c6 | 1,128 | 2.43 | Condition | 2 | 18.018 | 9.009 | 15.875 | 3.239e-04 | 5.048e-04 |
| Fib_c7 | 670 | 1.44 | Condition | 2 | 4.052 | 2.026 | 6.315 | 0.012 | 0.014 |

## References

1. Khan, M. S. et al. Global epidemiology of heart failure. Nat Rev Cardiol 21, 717–734 (2024).

2. Mann, D. L. & Bristow, M. R. Mechanisms and Models in Heart Failure. Circulation 111, 2837– 2849 (2005).

3. Hill, J. A. & Olson, E. N. Cardiac Plasticity. New England Journal of Medicine 358, 1370–1380 (2008).

4. Bozkurt, B. Contemporary pharmacological treatment and management of heart failure. Nat Rev Cardiol 21, 545–555 (2024).

5. Nguyen, A. H. et al. Medical Management and Device-Based Therapies in Chronic Heart Failure. Journal of the Society for Cardiovascular Angiography & Interventions 2, 101206 (2023).

6. Howard, T., Spilias, N., Bhattacharya, S., Harb, S. & Puri, R. Novel devices for managing heart failure. Current Opinion in Cardiology 37, 261 (2022).

7. Sun, Q., Karwi, Q. G., Wong, N. & Lopaschuk, G. D. Advances in myocardial energy metabolism: metabolic remodelling in heart failure and beyond. Cardiovasc Res 120, 1996–2016 (2024).

8. Ghazal, R., Wang, M., Liu, D., Tschumperlin, D. J. & Pereira, N. L. Cardiac Fibrosis in the Multi-Omics Era: Implications for Heart Failure. Circulation Research 136, 773–802 (2025).

9. Feng, H. et al. Macrophages in ventricular remodeling and heart failure: orchestrators of inflammation and repair. Front. Immunol. 16, (2025).

10. Fragasso, G. et al. The crosstalk between immune activation and metabolism in heart failure. A scientific statement of the Heart Failure Association of the ESC. European Journal of Heart Failure 27, 1700–1719 (2025).

11. Tan, S., Yang, J., Hu, S. & Lei, W. Cell-cell interactions in the heart: advanced cardiac models and omics technologies. Stem Cell Res Ther 15, 362 (2024).

12. Kyriakopoulos, C. P. et al. LVAD as a Bridge to Remission from Advanced Heart Failure: Current Data and Opportunities for Improvement. Journal of Clinical Medicine 11, 3542 (2022).

13. Topkara, V. K. et al. Recovery With Temporary Mechanical Circulatory Support While Waitlisted for Heart Transplantation. Journal of the American College of Cardiology 79, 900–913 (2022).

14. Lenneman, A. J. & Birks, E. J. Treatment Strategies for Myocardial Recovery in Heart Failure. Curr Treat Options Cardio Med 16, 287 (2014).

15. Tseliou, E. et al. Biology of myocardial recovery in advanced heart failure with long-term mechanical support. The Journal of Heart and Lung Transplantation 41, 1309–1323 (2022).

16. Ruppert, M. et al. Myocardial reverse remodeling after pressure unloading is associated with maintained cardiac mechanoenergetics in a rat model of left ventricular hypertrophy. American Journal of Physiology-Heart and Circulatory Physiology 311, H592–H603 (2016).

17. Rodrigues, P. G. et al. The Degree of Cardiac Remodelling before Overload Relief Triggers Different Transcriptome and miRome Signatures during Reverse Remodelling (RR)—Molecular Signature Differ with the Extent of RR. International Journal of Molecular Sciences 21, 9867 (2020).

18. Ramirez Flores, R. O., et al. Consensus Transcriptional Landscape of Human End-Stage Heart Failure. JAHA 10, e019667 (2021).

19. Amrute, J. M. et al. Defining cardiac functional recovery in end-stage heart failure at single-cell resolution. Nat Cardiovasc Res 2, 399–416 (2023).

20. Kuppe, C. et al. Spatial multi-omic map of human myocardial infarction. Nature 608, 766–777 (2022).

21. Lanzer, J. D. et al. A cross-study transcriptional patient map of heart failure defines conserved multicellular coordination in cardiac remodeling. Nat Commun 16, 9659 (2025).

22. Koenig, A. L. et al. Single-cell transcriptomics reveals cell-type-specific diversification in human heart failure. Nat Cardiovasc Res 1, 263–280 (2022).

23. Theall, B. & Alcaide, P. The heart under pressure: immune cells in fibrotic remodeling. Current Opinion in Physiology 25, 100484 (2022).

24. Liu, X. et al. Lymphoangiocrine signals promote cardiac growth and repair. Nature 588, 705–711 (2020).

25. Yang, Z. et al. Cross-talk between cardiac lymphatics and immune cells regulates inflammatory response and cardiac recovery after myocardial infarction. Front Immunol 16, 1557250 (2025).

26. Vieira, J. M. et al. The cardiac lymphatic system stimulates resolution of inflammation following myocardial infarction. J Clin Invest 128, 3402–3412 (2018).

27. Xu, C. et al. Automatic cell-type harmonization and integration across Human Cell Atlas datasets. Cell 186, 5876–5891.e20 (2023).

28. Drakos, S. G. et al. Impact of Mechanical Unloading on Microvasculature and Associated Central Remodeling Features of the Failing Human Heart. Journal of the American College of Cardiology 56, 382–391 (2010).

29. Simmonds, S. J. et al. Pericyte loss initiates microvascular dysfunction in the development of diastolic dysfunction. Eur Heart J Open 4, oead129 (2024).

30. Ramirez Flores, R., Lanzer, J., Dimitrov, D., Velten, B. & Saez-Rodriguez, J. Multicellular factor analysis of single-cell data for a tissue-centric understanding of disease. eLife 12, e93161 (2023).

31. Litviňuková, M. et al. Cells of the adult human heart. Nature 588, 466–472 (2020).

32. Dimitrov, D. et al. LIANA+ provides an all-in-one framework for cell-cell communication inference. Nat Cell Biol 26, 1613–1622 (2024).

33. Chaher, N. et al. Non-invasive in vivo imaging of changes in Collagen III turnover in myocardial fibrosis. npj Imaging 2, 33 (2024).

34. Shamhart, P. E. & Meszaros, J. G. Non-fibrillar collagens: Key mediators of post-infarction cardiac remodeling? Journal of Molecular and Cellular Cardiology 48, 530–537 (2010).

35. Williams, L. M. et al. Identifying collagen VI as a target of fibrotic diseases regulated by CREBBP/EP300. Proceedings of the National Academy of Sciences 117, 20753–20763 (2020).

36. Browaeys, R., Saelens, W. & Saeys, Y. NicheNet: modeling intercellular communication by linking ligands to target genes. Nat Methods 17, 159–162 (2020).

37. Marinescu, K. K., Uriel, N., Mann, D. L. & Burkhoff, D. Left ventricular assist device-induced reverse remodeling: it’s not just about myocardial recovery. Expert Rev Med Devices 14, 15–26 (2017).

38. Hilse, M. S. et al. Analysis of Metabolic Markers in Patients with Chronic Heart Failure before and after LVAD Implantation. Metabolites 11, 615 (2021).

39. Biggs, R. M. et al. Persistent Fibrosis in Heart Failure With a Reduced Ejection Fraction Linked to Phenotypic Differences in Human Cardiac Fibroblast Populations. J Am Heart Assoc 14, e039747 (2025).

40. Cheng, L. et al. Long-Term Follow-Up of the TRED-HF Trial: Implications for Therapy in Patients with Dilated Cardiomyopathy and Heart Failure Remission. Eur J Heart Fail 27, 113–123 (2025).

41. Roth, R. et al. Transcriptional signature of cardiac myocyte recovery in mice and human reveals persistent upregulation of epigenetic factors. Epigenetics 20, 2506625.

42. Halliday, B. P. et al. Withdrawal of pharmacological treatment for heart failure in patients with recovered dilated cardiomyopathy (TRED-HF): an open-label, pilot, randomised trial. The Lancet 393, 61–73 (2019).

43. Li, X. et al. Ventricular assist device unloading reverses microvascular senescence in single ventricle disease. Nat Cardiovasc Res 5, 262–280 (2026).

44. Farbehi, N. et al. Single-cell expression profiling reveals dynamic flux of cardiac stromal, vascular and immune cells in health and injury. Elife 8, e43882 (2019).

45. Kattih, B. et al. Single-nuclear transcriptome profiling identifies persistent fibroblast activation in hypertrophic and failing human hearts of patients with longstanding disease. Cardiovasc Res 119, 2550–2562 (2023).

46. Daseke, M. J., Tenkorang, M. A. A., Chalise, U., Konfrst, S. R. & Lindsey, M. L. Cardiac fibroblast activation during myocardial infarction wound healing: Fibroblast polarization after MI. Matrix Biol 91–92, 109–116 (2020).

47. Peisker, F. et al. Mapping the cardiac vascular niche in heart failure. Nat Commun 13, 3027 (2022).

48. Komuro, J. et al. Heart failure-specific cardiac fibroblasts contribute to cardiac dysfunction via the MYC–CXCL1–CXCR2 axis. Nat Cardiovasc Res 4, 1135–1151 (2025).

49. Kumari, M. et al. Activation of IRF3 in cardiomyocytes impairs mitochondrial oxidative function through PGC-1α inhibition and drives heart failure. Nat Commun 17, 2051 (2026).

50. Melika, E. T. et al. Interferon gamma signaling drives cardiac metabolic rewiring. 2025.08.31.673391 Preprint at 10.1101/2025.08.31.673391 (2025).

51. Beffert, U. et al. Functional dissection of Reelin signaling by site-directed disruption of Disabled-1 adaptor binding to apolipoprotein E receptor 2: distinct roles in development and synaptic plasticity. J Neurosci 26, 2041–2052 (2006).

52. Bock, H. H. & May, P. Canonical and Non-canonical Reelin Signaling. Front Cell Neurosci 10, 166 (2016).

53. Alexander, A., Herz, J. & Calvier, L. Reelin through the years: From brain development to inflammation. Cell Reports 42, 112669 (2023).

54. Ding, Y. et al. Loss of Reelin protects against atherosclerosis by reducing leukocyte-endothelial cell adhesion and lesion macrophage accumulation. Sci Signal 9, ra29 (2016).

55. Pei, L. et al. Thrombospondin 1 and Reelin act through Vldlr to regulate cardiac growth and repair. Basic Res Cardiol 119, 169–192 (2024).

56. Kashihara, T. et al. YAP mediates compensatory cardiac hypertrophy through aerobic glycolysis in response to pressure overload. J Clin Invest 132, e150595 (2022).

57. Ninh, V. K. et al. Cardiomyocyte YAP represses myocardial inflammation and fibrosis and restrains MEF2-regulated gene expression. Am J Physiol Heart Circ Physiol 329, H774–H787 (2025).

58. Jover, E. et al. Expression of the lymphangiogenic reelin is associated with sex-dependent calcific aortic stenosis in men. Atherosclerosis 403, 119162 (2025).

59. Melleby, A. O. et al. A novel method for high precision aortic constriction that allows for generation of specific cardiac phenotypes in mice. Cardiovasc Res 114, 1680–1690 (2018).

60. Burridge, P. W. et al. Chemically defined generation of human cardiomyocytes. Nat Methods 11, 855–860 (2014).

61. D’Arcangelo, G. et al. Reelin Is a Secreted Glycoprotein Recognized by the CR-50 Monoclonal Antibody. J. Neurosci. 17, 23–31 (1997).

62. Love, M., Huber, W. & Anders, S. Moderated estimation of fold change and dispersion for RNA-seq data with DESeq2. Genome Biol 15, 550 (2014).

63. Zhu, A., Ibrahim, J. G. & Love, M. I. Heavy-tailed prior distributions for sequence count data: removing the noise and preserving large differences. Bioinformatics 35, 2084–2092 (2019).

64. Badia-I-Mompel, P. et al. decoupleR: ensemble of computational methods to infer biological activities from omics data. Bioinformatics Advances 2, vbac016 (2022).

65. Müller-Dott, S. et al. Expanding the coverage of regulons from high-confidence prior knowledge for accurate estimation of transcription factor activities. Nucleic Acids Res 51, 10934–10949 (2023).

66. Liberzon, A. et al. The Molecular Signatures Database (MSigDB) hallmark gene set collection. Cell Syst 1, 417–425 (2015).

67. Wolf, F., Angerer, P. & Theis, F. SCANPY: large-scale single-cell gene expression data analysis. Genome Biol 19, 15 (2018).

68. Wolock, S. L., Lopez, R. & Klein, A. M. Scrublet: Computational Identification of Cell Doublets in Single-Cell Transcriptomic Data. Cell Systems 8, 281–291.e9 (2019).

69. Korsunsky, I. et al. Fast, sensitive and accurate integration of single-cell data with Harmony. Nat Methods 16, 1289–1296 (2019).

70. Kanemaru, K. et al. Spatially resolved multiomics of human cardiac niches. Nature 619, 801–810 (2023).

71. Tucker, N. R. et al. Transcriptional and Cellular Diversity of the Human Heart. Circulation 142, 466–482 (2020).

72. Tirosh, I. et al. Dissecting the multicellular ecosystem of metastatic melanoma by single-cell RNA-seq. Science 352, 189–196 (2016).

73. Robinson, M. D., McCarthy, D. J. & Smyth, G. K. edgeR: a Bioconductor package for differential expression analysis of digital gene expression data. Bioinformatics 26, 139–140 (2010).

74. Argelaguet, R. et al. MOFA+: a statistical framework for comprehensive integration of multi-modal single-cell data. Genome Biology 21, 111 (2020).

75. Türei, D. et al. OmniPath: integrated knowledgebase for multi-omics analysis. Nucleic Acids Res 54, D652–D660 (2026).

76. Shannon, P. et al. Cytoscape: A Software Environment for Integrated Models of Biomolecular Interaction Networks. Genome Res. 13, 2498–2504 (2003).

